# The neural computation of human prosocial choices in complex motivational states

**DOI:** 10.1101/851931

**Authors:** Anne Saulin, Ulrike Horn, Martin Lotze, Jochen Kaiser, Grit Hein

## Abstract

Motives motivate human behavior. Most behaviors are driven by more than one motive, yet it is unclear how different motives interact and how such motive combinations affect the neural computation of the behaviors they drive. To answer this question, we induced two prosocial motives simultaneously (multi-motive condition) and separately (single motive conditions). After the different motive inductions, participants performed the same choice task in which they allocated points in favor of the other person (prosocial choice) or in favor of themselves (egoistic choice). We used fMRI to assess prosocial choice-related brain responses and drift diffusion modelling to specify how motive combinations affect individual components of the choice process. Our results showed that the combination of the two motives in the multi-motive condition increased participants’ choice biases prior to the behavior itself. On the neural level, these changes in initial prosocial bias were associated with neural responses in the bilateral dorsal striatum. In contrast, the efficiency of the prosocial decision process was comparable between the multi-motive and the single-motive conditions. These findings provide insights into the computation of prosocial choices in complex motivational states, the motivational setting that drives most human behaviors.

**Highlights:**

- Activating different social motives simultaneously can enhance prosocial choices
- Multi-motive combinations change initial prosocial biases
- Dorso-striatal activation increases with larger increase of prosocial bias
- Multi-motive combinations modulate relative response caution

## 1. Introduction

All choice behaviors are incited by motives, which can be complex. Documenting this motivational complexity, many animal (Jennings et al., 2013; Kennedy and Shapiro, 2009) and most human behaviors are driven by multiple motives that are active at the same time, and affect each other (Engel and Zhurakhovska, 2016; Hughes and Zaki, 2015; Jagers et al., 2017; Kruglanski et al., 2018; Lewin et al., 1951; Takeuchi et al., 2015; Terlecki and Buckner, 2015). For example, the decision to help an elderly relative is often driven by empathy with her needs, and at the same time, by the wish to reciprocate help received by this person in the past, i.e, the social norm of reciprocity. Consequently, most choice behaviors are driven by combinations of different motives and cannot be explained by one “motivational force” alone. However, the combination of motives is not directly observable. Thus, to understand and predict choice behaviors, it is crucial to elucidate the neuro-computational mechanisms through which multiple simultaneously activated motives affect behavioral choice processes.

The processing of single-motive states and its impact on behavioral choices in animals (e.g., place preferences) (Jennings et al., 2013) have been linked to dopaminergic neurons in the striatum (Kim and Im, 2018; Robinson et al., 2006; Salamone and Correa, 2012). In line with these results, human neuroscience studies have shown that the striatum is involved in the processing of different individual motives, as well as motivated choice behaviors, both in the social (Báez-Mendoza and Schultz, 2013; Bhanji and Delgado, 2014) and non-social domain (Salamone et al., 2016; Shohamy, 2011). In more detail, the ventral striatum has been linked to the learning and encoding of values and the predictions of future rewards (Kable and Glimcher, 2007; Liljeholm and O ’Doherty, 2012; O’Doherty et al., 2004; Strait et al., 2015), whereas the dorsal striatum has been linked to initiating and optimizing choices based on these encoded values (Balleine et al., 2007; Liljeholm and O ’Doherty, 2012; O’Doherty et al., 2004; Palmiter, 2008; Robinson et al., 2006).Together, this previous work has provided insights into the neural underpinnings of individual motivational processes. However, the neural computation of behaviors that are driven by different motives remains unclear.

To address this issue, we developed a paradigm in which participants made the same choices (prosocial vs. egoistic) based on different, simultaneously activated motives (multi-motive condition), or based on each of these motives separately (single-motive conditions). Specifically, we studied the effect of simultaneously activated social motives in a social choice paradigm in which participants repeatedly had the choice between a prosocial and an egoistic option (**Fig. 1A**). Inspired by an influential model of prosocial motivations (Batson et al., 2011), we induced two key motives that both incite prosocial behavior - the empathy motive, defined as the affective response to another person’s misfortune (Batson et al., 1995; Hein et al., 2016a; Lamm et al., 2011), and the reciprocity motive, defined as the desire to reciprocate perceived kindness with a kind behavior (Gouldner, 1960; Hein et al., 2016a; McCabe et al., 2003).

**Figure 1.**
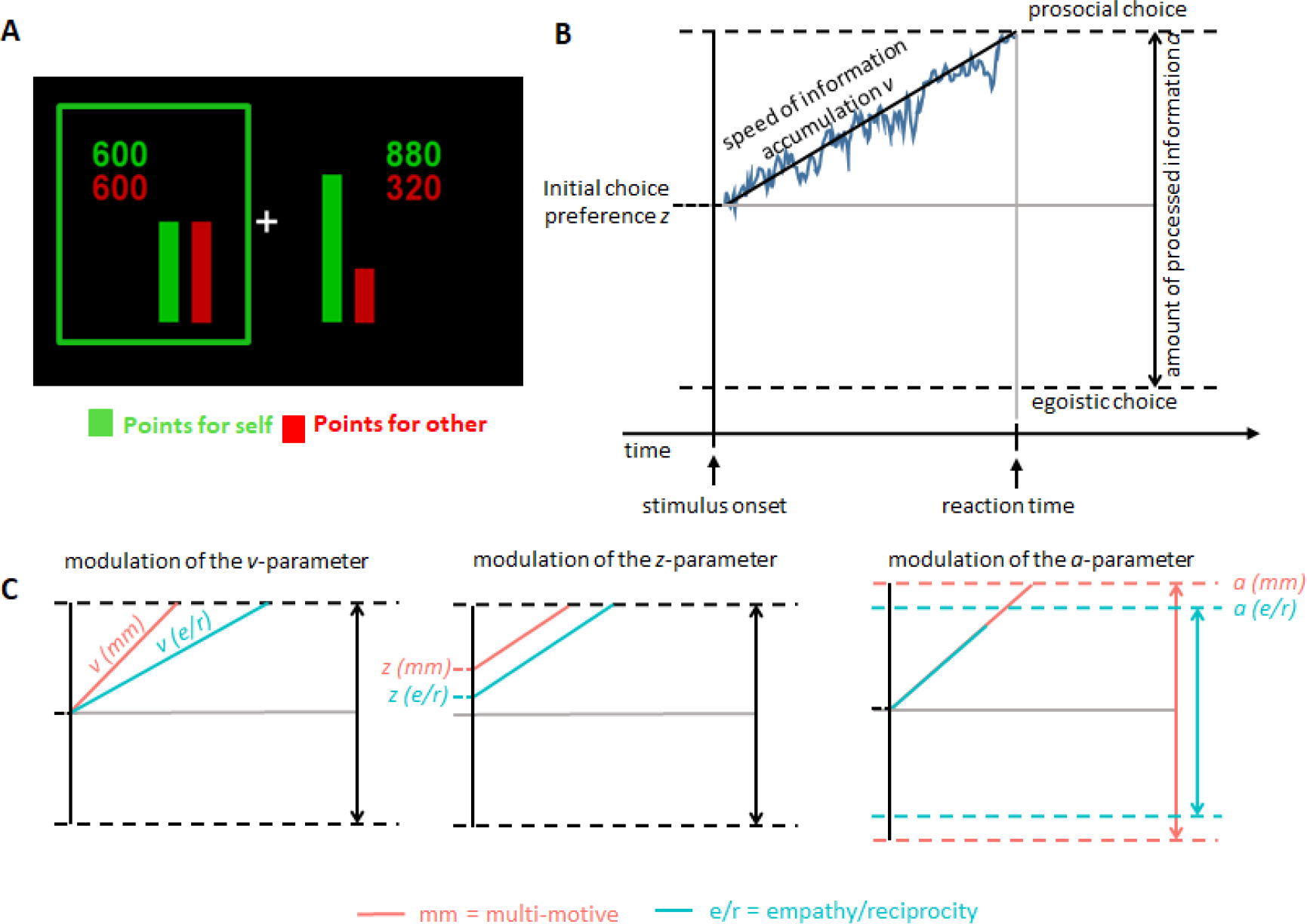
Example of point allocation during the choice task, schematic illustration of the drift-diffusion model and hypotheses regarding the impact of different drift-diffusion parameters on the choice process in multi-motive and single-motive conditions. **(A)** Participants chose between a prosocial and an egoistic option to allocate points to themselves (in this example shown in green) and a partner (in this example shown in red). Colors were counter-balanced across participants. In this example trial, the participant chose the prosocial option, which maximized the outcome of the partner at a cost to the participant (green box). **(B)** The drift-diffusion model conceptualizes the choice process as noisy accumulation of information (squiggly blue line). The *v*-parameter describes the speed at which information is accumulated in order to choose one of the options, i.e., the efficiency of the choice process itself. The *z*-parameter reflects the initial choice bias, i.e., the degree to which an individual prefers one of the choice options prior to making the choice. The third component, parameter *a*, quantifies the amount of relative evidence that is required to choose one of the options. Once the accumulated information reaches either boundary, the choice is made (upper boundary = prosocial choice; lower boundary = egoistic choice). **(C)** An enhancement of prosocial choice frequency in the multi-motive condition (red) compared to the single motive conditions (i.e., the empathy or the reciprocity condition; blue) may result from an increased speed of information accumulation (*v*-parameter; left panel), and/or an increased initial bias toward making a prosocial choice (*z*-parameter; middle panel). On average, the amount of required relative evidence (*a*-parameter) may be higher in the multi-motive condition compared to the single motive conditions (right panel).

In combination with fMRI and hierarchical drift-diffusion modeling (hierarchical DDM) (Forstmann et al., 2016; Ratcliff et al., 2016; Vandekerckhove et al., 2011; Wiecki et al., 2013), this paradigm allowed us to specify how the combination of different motives affects individual components of neural goal-directed (i.e., prosocial) choice computation, compared to computation of the same choice in a simple motivational state (i.e., driven by only one of the two motives).

Drift-diffusion models (DDMs) characterize how noisy information is accumulated to select a choice option (**Fig. 1B**) based on three different parameters (the *v*, *z* and *a* parameters) (Forstmann et al., 2016; Ratcliff et al., 2016). The *v*-parameter describes the speed at which information is accumulated in order to choose one of the options, i.e., the efficiency of the choice process itself. The *z*-parameter reflects the initial choice bias, i.e., the degree to which an individual prefers one of the choice options prior to making the choice. Thus, in contrast to the *v*-parameter, which models the choice process itself, the *z*-parameter models the individual bias with which a person enters the choice process. For example, if a person has a strong initial bias towards prosocial choices (reflected by a large value of the parameter *z*), the starting point of the choice computation is located closer to the prosocial choice boundary, and thus, this person is more likely to choose the prosocial option. The third component, parameter *a*, quantifies the amount of relative evidence that is required to choose one of the options.

Previous neuroscience studies have identified brain regions that are associated with changes of these choice parameters. For example, it has been shown that reward-related improvement of perceptual discrimination is driven by changes in the *z*-parameter, related to changes of frontoparietal activation (Mulder et al., 2012). Another study using a task from the perceptual domain has shown that increased evidence accumulation under time pressure is linked to increased activation in premotor regions (preSMA) and the dorsal striatum (Forstmann et al., 2008). Other studies have used similar modelling approaches to investigate value-based decisions (Gluth et al., 2012; Hare et al., 2011), for example using a buying task in which participants could decide to accept or reject a stock after receiving probabilistic information about the stock from different rating companies (Gluth et al., 2012). As a main result, Gluth and colleagues showed that the amount of relative evidence that participants required for making a choice was related to the neural response in the anterior insula (AI) and the dorsal striatum. The finding in dorsal striatum resembled evidence from DDM studies obtained with perception paradigms (Forstmann et al., 2008) and indicates that, the striatum is a plausible neural candidate for tracking changes in choice components in different motivational settings (e.g., induced by time pressure, Forstmann et al., 2008, or by others’ information, Gluth et al., 2012).

In our study, we modeled the three relevant choice parameters (*v, z*, and *a*) for choices that were driven by the combination of the two motives and for the same choices that were driven by each of the motives separately. It is important to note that the choice process may also be influenced by other motives than empathy and reciprocity, i.e., the motives that were experimentally induced. That said, our paradigm can provide insights into the multi-motive choice process even if other motives are potentially activated, because multi-motive choices are contrasted with the same choices that are driven by the respective single motives.

According to one hypothesis, the simultaneous activation of multiple motives may facilitate the computation of the choice option that is favored by the motives. In the present paradigm this means that computation of the prosocial choice option should be facilitated since empathy and reciprocity both drive prosocial behavior. In this case, we should observe an increase in prosocial behavior in the multi-motive condition (empathy and reciprocity motive active) compared to the single-motive conditions (only empathy or only reciprocity active) that cannot be explained by the difference between the single-motive conditions. Specifying the mechanism underlying such a facilitation, the DDM proposes that a facilitation of prosocial choices in the multi-motive condition may originate A) from an increased speed of information accumulation (*v*-parameter; **Fig. 1C**, left panel (Flagan et al., 2017; Janczyk and Lerche, 2019; Krajbich et al., 2015)), B) from an enhancement of participants’ initial bias to choose the prosocial option (*z*-parameter; **Fig. 1C**, middle panel; (Chen and Krajbich, 2018; Mulder et al., 2012; Toelch et al., 2018), or C) from an enhancement of the *v*-as well as the *z*-parameter in the multi-motive condition, compared to the single-motive condition.

Alternatively, we may observe fewer prosocial decisions in the multi-motive condition compared to the single motive conditions, reflected by a decreased speed of information accumulation (lower DDM *v*-parameter) and/or decreased initial bias to choose the prosocial option (lower DDM *z*-parameter). Moreover, in the multi-motive condition, participants are required to process two motives simultaneously, in addition to the trial-by-trial choice option information (which was constant across all conditions because participants performed the identical choice task). This additional motive-related information may cause participants to make more careful responses in the multi-motive condition and thus increase the *a*-parameter in the multi-motive condition compared to the single-motive conditions (**Fig.1C**, right panel).

Regarding the underlying neural mechanisms, we hypothesized that changes in DDM choice parameters in the multi-motive compared to the single-motive conditions might be related to changes of activation in the ventral striatum, i.e., a region that is involved in the integration of different choice values (here the value of empathy-based and the value of reciprocity-based choices; (Kable and Glimcher, 2007; Liljeholm and O ’Doherty, 2012; O’Doherty et al., 2004; Strait et al., 2015), and/ or activation of the dorsal striatum, i.e., a region that is related to integration of choice preferences that derive from these different choice values (Balleine et al., 2007; Liljeholm and O ’Doherty, 2012; O’Doherty et al., 2004; Palmiter, 2008; Robinson et al., 2006).

## 2. Material and methods

### 2.1. Participant details

Forty-two right-handed healthy female participants (mean age = 23.1 years, SD = 2.8 years) and four female confederates took part in the experiment. We chose female participants as well as female confederates in order to control for gender and avoid cross-gender effects. The confederates were students who had been trained to serve in all the different conditions counterbalanced across participants. Prior to the experiment, written informed consent was obtained from all the participants. The study was approved by the local ethics committee (BB 023/17). Participants received monetary compensation. Three participants had to be excluded due to technical problems and dropout. Another subject had to be excluded due to excessive head movements (more than 5% of the scans contained rapid head motion with more than 0.5 mm displacement per TR). Five participants had to be excluded as outlier based on their choices (less than ten prosocial choices across all condition; three standard deviations above the mean in central measures). Thus, we analyzed 33 data sets using a within-subjects design. We aimed for 34 data sets, which corresponds to the median sample size of neuroimaging studies determined in a recent review (N = 33; Yeung, 2018). We tested 40 participants to meet this target, accounting for a drop-out rate of about 15% which, based on our experience, is common in fMRI studies. Our final analyses includes 33 data sets, in accordance with the median sample reported by Yeung (2018). Given that it is difficult to collect large data sets with expensive and time-consuming methods like fMRI, the importance of stringent statistical thresholds is highlighted (Carter et al., 2016; Roiser et al., 2016; Woo et al., 2014; Yeung, 2018). To analyse the results of the second level regression we thus used cluster-level family wise error correction at the whole brain level after applying a threshold of p < .001 on an uncorrected level. Neural activations that are thresholded at this level are seen as valid and reliable (Eklund et al., 2016; Woo et al., 2014; Yeung, 2018).

### 2.2. Procedure

Prior to the motive induction and choice task, the individual thresholds for pain stimulation were determined for the participants and all the confederates (see section 2.5. Pain stimulation for details). Next, the participants and confederates were assigned their different roles by a manipulated lottery (drawing matches). In order to ensure that each participant was always assigned her designated role as a participant (pain recipient during motive induction; decider during the decision task), the drawing of the matches was organized in such a way that she always drew the last match. The confederates were assigned the roles of the empathy partner, the reciprocity partner, the multi-motive partner or the baseline partner, and these roles were counterbalanced across participants. In accordance with these roles, two of the confederates first went to an ostensible other experiment and the other two waited to be seated in the scanner room. Each confederate was matched with a specific color and seating position (to the left vs. to the right of the fMRI scanner), and their color designation and seating positions were counter-balanced across participants. Next, the first two confederates (the empathy partner, reciprocity partner, multi-motive partner, or baseline partner) were seated to the left and the right of the participant who was lying inside the fMRI scanner and the first motive induction took place (for overview of an example procedure, see **Fig. 2**). After the motive induction, image acquisition for the choice task was started, during which the participant allocated points to her respective partners. This way the participant only had to remember interactions with two partners at any one time. After the choice task, the first confederates were replaced by the other two confederates and the second part commenced. Part 2 had the same structure as part 1: first, the participant underwent motive induction 2 followed again by the choice task. The order of motive inductions and the type of partner the confederates represented were counterbalanced across participants.

**Figure 2.**
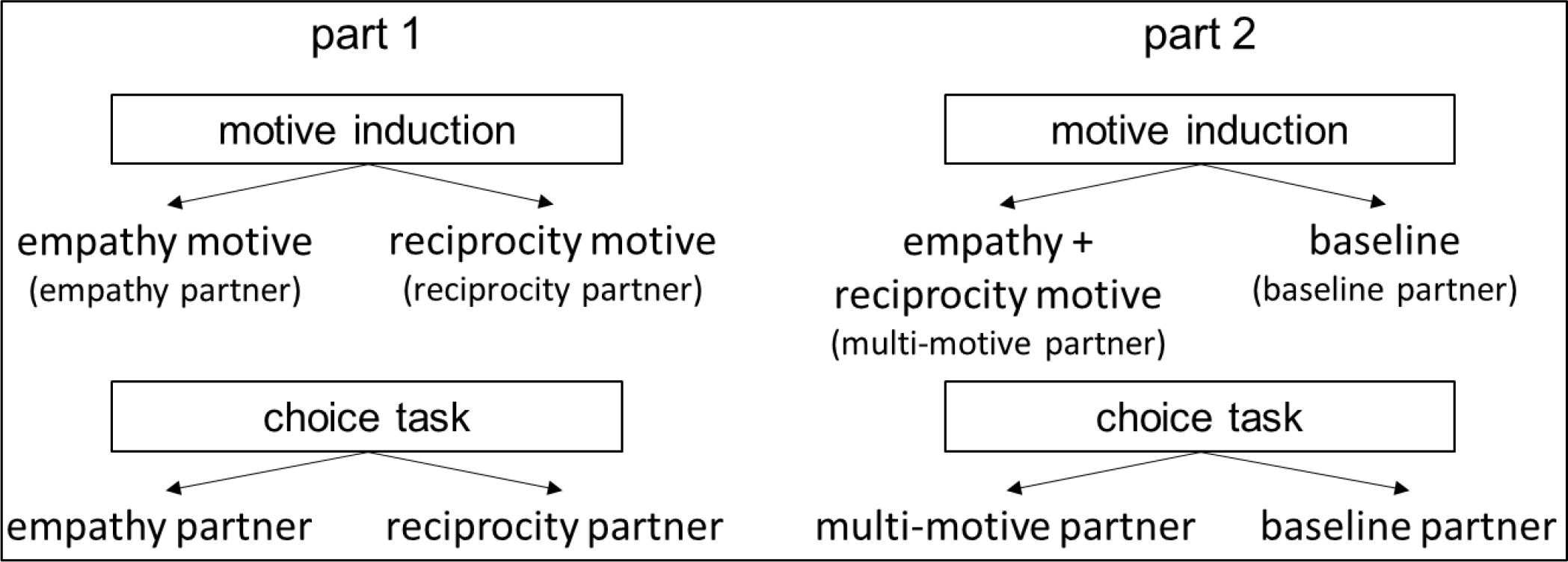
Overview of an example experimental procedure. The study consisted of two parts. In this example, in part 1, the empathy motive was activated towards one confederate (the empathy partner) and the reciprocity motive was activated towards the other confederate (the reciprocity partner). In the following choice task, participants allocated points to the empathy partner (i.e., driven by the empathy motive) or the reciprocity partner (i.e., driven by the reciprocity motive). Next, the confederates were replaced by two new individuals that served as partners for part 2. In part 2, the empathy and the reciprocity motive were activated simultaneously towards one confederate (multi-motive partner) and no motive was induced towards the other confederate (baseline partner). In the following choice task, participants allocated points towards the multi-motive partner (i.e., driven by two motive simultaneously) and towards the baseline partner (i.e., independently of any motive induction). The order of motive induction (empathy, reciprocity, multi-motive, baseline) was counterbalanced across participants and the four confederates. The respective partner was indicated by a cue in one of four counterbalanced colors.

At the end of the experiment, all the confederates left and the participant remained in the scanner until anatomical image acquisition was completed. Finally, participants were asked to complete a questionnaire measuring trait aspects of empathy (IRI, Davis, 1983; Jordan et al., 2016) and reciprocity (Perugini et al., 2003). Participants spent approximately 60 min in the scanner and the entire procedure lasted approximately 2.5 hours. To avoid possible reputation effects, which could influence participants’ behavior, participants were informed that they would not meet the

### 2.3. Motive inductions

#### 2.3.1. Empathy induction

During the study, participants were paired with four partners (confederates of the experimenter). Participants saw the hand of the respective partners with the attached pain electrode. In the empathy condition, the participants repeatedly observed one of the confederates (the empathy partner) receiving painful shocks in a number of trials, a situation known to elicit an empathic response (Batson et al., 1995; Hein et al., 2016a; Lamm et al., 2011). Each empathy-induction trial started with a colored arrow shown for 1,000 ms, which indicated the empathy partner. After this cue and a jittered (1,000–2,000 ms) fixation cross, the same colored flash was displayed for 1,500 ms. Participants were informed that a dark-colored flash indicated that the corresponding partner received a painful stimulus at that moment; a light-colored flash indicated a non-painful stimulus. During (ostensible) stimulation of the respective partner, participants either saw a dark colored flash (painful stimulation) or a light colored flash (non-painful stimulation). Since all partners were confederates of the experimenter, they did not actually receive painful stimulations. Thus, the trials in which participants saw the dark colored flash were “ostensibly painful” for the partner. To assess the success of the empathy induction, a rating scale was shown for a maximum of 6 s. Participants reported how they felt after observing the partner receive painful or non-painful stimuli (“How do you feel?” in German). The scale ranged from -4 (labeled “very bad”) to +4 (labeled “very good”) with intervals of one and was visually displayed. Before analysis, the induction ratings were recoded such that high positive values reflect strong responses to the induction procedure (strong empathy motive). Participants had to respond within 6 s. The inter-trial interval was 1,500 ms. Empathy induction consisted of 12 trials: nine of which were ostensibly painful for their partner (i.e., the confederate).

#### 2.3.2. Reciprocity induction

The reciprocity motive is defined as the desire to reciprocate perceived kindness with kind behavior (Gouldner, 1960; Hein et al., 2016a; McCabe et al., 2003). Therefore, in the reciprocity condition, we induced the reciprocity motive by instructing one of the confederates (the reciprocity partner) to give up money in several trials to save the participant from painful shocks (Hein et al., 2016a). Each reciprocity-induction trial also started with an arrow colored in the reciprocity partner’s color, which pointed toward the seating position of the reciprocity partner (left or right) and was shown for 1,000 ms. Next, the participants were shown a flash displayed to the right and a crossed-out flash displayed to the left of a centered fixation cross. Participants were told that this was the decision screen, which the reciprocity partner also saw while making her decision to either save or not save the participant from painful stimulation. After a jittered interval of 2,000 to 4,000 ms, a box appeared around one of the flashes, indicating the ostensible choice of the reciprocity partner. Depending on where the box was displayed, the reciprocity partner had either decided to forego a monetary award of 2 € in order to save the participant from painful stimulation (a box around the crossed-out flash) or decided to take the money and not save the participant (a box around the flash that was not crossed-out). After 1000ms participants rated how they felt about the decision of the partner (“How do you feel?” in German). The ratings were recoded such that high positive values reflect a strong positive response to the decision of the partner, indicating a strong reciprocity motive. After a jittered (1,000 to 2,000 ms) fixation cross, the participant saw an information on the screen, indicating whether the decision of the reciprocity partner would be implemented (“decision accepted”) or not (“decision declined”), displayed for 1000 ms. This additional stage was included in order to ensure the same amount of painful stimulations were administered across all conditions (50 %), while at the same time allowing for the high rate (75 %; 9 out of 12 trials) of the reciprocity partner’s decisions to help. While instructing the participants, it was highlighted that the choice of the reciprocity partner reflected her willingness (or unwillingness) to help, while a computer algorithm decided about the implementation of the decision.

Thus, four types of reciprocity trials were possible. When the partner decided to save the participant from painful stimulation and this decision was accepted, the participant did not receive a painful stimulus, which was visually represented by a crossed-out flash (1,500 ms). However, when the reciprocity partner’s decision to save the participant was declined, participants received a painful stimulus, which was accompanied by the display of a flash (1,500 ms). Similarly, when the partner decided not to save the participant and this decision was accepted, the participant received a painful stimulus accompanied by the display of a flash. Finally, when the partner decided to not save the participant and this decision was declined, the participant did not receive painful stimulation, which was visually represented by a crossed-out flash. The inter trial fixation cross was displayed for 1,500 ms before the next trial started.

#### 2.3.3. Multi-motive induction

In the multi-motive condition, the participants repeatedly observed how one of the confederates (the multi-motive partner) received painful shocks and also gave up money to spare the participant from painful shocks. The multi-motive induction procedure combined the empathy- and reciprocity-induction procedures. As in the empathy-induction condition, it included 12 empathy induction trials, nine of which were ostensibly painful for the partner. As in the reciprocity-induction condition, it included 12 reciprocity trials, of which participants received help in nine out of 12 trials. The stimulation and trial structure were identical to the empathy- and reciprocity-induction trials described above, except that the relevant colors were replaced by the colors matched to the multi-motive partner (i.e., the color of the pain flash in the empathy trials and the color of the box highlighting the decision of the partner in the reciprocity trials).

#### 2.3.4. Additional control trials for empathy and reciprocity induction

In order to equalize the number and types of trials (i.e., the length and structure of the interaction with each motive partner) across conditions, the empathy-induction procedure also included trials that were identical to the reciprocity trials, except that the computer decided whether the participant would be saved from a painful stimulus and not the empathy partner. This computer’s decision was visually represented by a white-colored box appearing either around the crossed-out flash (saving the participant) or the normal flash (not saving the participant). It was clearly explained to each participant that the color white was not matched with any of the partners but indicated the computer’s choice. The empathy-induction procedure consisted of 12 control trials, in addition to the 12 empathy trials described above, resulting in 24 trials, i.e., the identical number of trials as the multi-motive induction procedure. Similarly, the reciprocity-induction procedure included trials that were identical to the empathy-induction trials, except that the reciprocity partner only received non-painful stimulation on these trials, as visually represented by a light-colored flash. In total, the reciprocity-induction procedure consisted of 12 of these control trials and 12 reciprocity trials (see above), i.e., 24 trials (identical to the other conditions).

#### 2.3.5. Baseline induction

The baseline procedure consisted of 24 trials in total, 12 trials in which the baseline partner only received non-painful stimulation and 12 trials in which the computer decided whether the participant would be saved from a painful stimulus or not. This computer’s decision was visually represented by a white-colored box either appearing around the crossed-out flash (saving the participant) or the normal flash (not saving the participant). It was clearly explained to the participant that the white box did not represent the decision of a person but indicated the computer’s choice.

### 2.4. Choice task

After the motive inductions, participants performed a social choice task inside the fMRI scanner. The choice task was a two-alternative-forced-choice adaptation of the commonly used Dictator Game (Forsythe et al., 1994), which has been successfully used in previous studies (e.g., Chen and Krajbich, 2018; Hein et al., 2016b; Krajbich et al., 2015). In each trial of this choice task, participants allocated money to themselves and one of the partners (**Fig. 1A**) and could choose between maximizing the relative outcome of the other person by reducing their own relative outcome (prosocial choice) and maximizing their own relative outcome at a cost to the partner (egoistic choice). The outcome was relative to the outcome that the participant would have gained when choosing the other option. The initial number of points was always higher for the participant compared to the partners. This measure was inspired by previous behavioral economics research, showing that prosocial behaviors depend on the initial payoff allocation between the participant and the participant’s partner (Bolton and Ockenfels, 2000; Charness and Rabin, 2002; Fehr and Schmidt, 1999). In particular, if subjects have a lower initial payoff than their partner (“disadvantageous initial inequality”), they are much less willing to behave altruistically toward the partner compared to a situation with advantageous initial inequality (i.e., when the participant has a higher initial payoff than the partner). The choice options used in the present study created advantageous inequality to optimize the number of prosocial choices, which was the main focus of our study. The exact point distributions are provided in **Table A1**.

Depending on the type of partner the participants faced in the choice task, there were four conditions – the empathy condition, the reciprocity condition, the multi-motive condition, and the baseline condition. Importantly, the choice task was identical in all the conditions.

In more detail, participants were asked repeatedly to choose between two different distributions of points that each represented different amounts of monetary pay-offs for themselves and one of the partners (see **Fig. 1A**). Each choice-trial started with a colored arrow shown for 1,000 ms, indicating the next interaction partner. After this cue, participants saw the two possible distributions of points in different colors, indicating the potential gain for the participant or for the current partner. Colors were counterbalanced across participants. Participants had to choose one of the distributions within 4,000 ms. A green box appeared around the distribution that was selected by the participant at 4,000 ms after distribution onset. The box was shown for 1,000 ms. The length of the inter-trial interval, as indicated by a fixation cross, was jittered between 4,000 and 6,000 ms. At the end of the experiment, two of the distributions chosen by the participant were randomly selected for payment (100 points = 50 cents). We analyzed 38 choice trials in each motive-induction condition, i.e., 152 trials in total. In addition to the 38 trials, each condition contained four trials in which the same choice option maximized the outcome of the participant and the partner (non-competitive trials). These trials were included to increase the variability of choices and thus to keep the participants engaged. They were excluded from the analyses because they could not be classified as prosocial or egoistic choice trial. For each condition and participant, the same distributions were used and presented in random order.

### 2.5. Pain stimulation

For pain stimulation, we used a mechano-tactile stimulus generated by a small plastic cylinder (513 g). The projectile was shot against the cuticle of the left index finger using air pressure (Impact Stimulator, Labortechnik Franken, Release 1.0.0.34). The criterion for painful stimulation was a subjective value of 8 on a pain scale ranging from 1 (no pain at all, but a participant could feel a slight touch of the projectile) to 10 (extreme, hardly bearable pain). The participants were told that a value of 8 corresponded to a painful, but bearable stimulus, and a non-painful stimulus corresponded to a value of 1 on the same subjective pain scale. These subjective pain thresholds were determined using a stepwise increase of air pressure (stepsize of 0.25 mg/s), starting with the lowest possible pressure (0.25 mg/s), which caused the projectile to barely touch the cuticle, and increasing in stimulus intensity until it reached a level that corresponded to the individual’s value of 8 (range = 2.75– 3.5 mg/s).

### 2.6. Experimental design and statistical analyses

#### 2.6.1. Regression analyses

Regression analyses were conducted using the R-packages “lme4 and “car” (R Core-Team, 2018). For mixed models, we report the chi-square values derived from Wald chisquare tests using the “Anova” (car package) function. For predefined contrasts we report the t-values derived from the summary() function. When more than one predictor was included in the model, the function emmeans was used in order to compute contrasts between factor levels.

To test the differences in induction ratings and the relationship between induction ratings and frequencies of prosocial choice, the mean induction ratings and frequencies of prosocial choices were calculated for each participant for each condition (empathy, reciprocity, multi-motive, and baseline) and entered as a dependent variable into mixed models with conditions (empathy, reciprocity, multi-motive, and baseline) and induction ratings as fixed effects and participants as random effects. Additionally, in order to probe the specificity of the induction procedure, we tested whether trait empathy (empathic concern subscale of the IRI, Davis (1983)) and trait reciprocity (PNR, (Perugini et al., 2003)) differentially influenced choice behavior in the three motive conditions (empathy, reciprocity, multi-motive). Specifically, we conducted a linear mixed model regression with the frequency of prosocial choices as dependent variable, trait measure type (empathy / reciprocity), individual trait measure scores (empathy / reciprocity), motive induction condition (empathy, reciprocity, multi-motive), and their interactions as fixed effects, and participant as random intercept.

To test whether prosocial behavior was influenced by trial-by-trial point information, condition and their interaction, a logistic mixed model regression was conducted with the possible gain for the partner (i.e. difference in points between the two options for the partner, |partner’s gain option 1 – partner’s gain option 2|), the possible loss for the participant (i.e., the difference in points between the two options for the participant, |participant’s gain option 1 – participant’s gain option 2|) and condition as predictor variables. The binary choice outcome (prosocial vs. egoistic choice) was used as dependent variable. To investigate the differences in prosocial behavior between the social motives (multi-motive > reciprocity and multi-motive > empathy) more closely, contrasts were calculated using the emmeans function.

To specifically test whether prosocial behavior was differentially influenced by inequity aversion, a logistic mixed model regression was conducted with the predictor variables condition and the difference in point equality of the participant’s and the partner’s outcome between the two choice options. To compute this variable, we first calculated calculating the difference between the gains for each option (i.e., |partner’s gain option 1 – participant’s gain option 1| for each choice option). Second, these differences were subtracted from each other in order to obtain a measure of point equality for each choice trial. Again, the binary choice outcome (prosocial vs. egoistic choice) was used as dependent variable.

To test whether the frequency of prosocial choices and reaction times were equally distributed across conditions, we conducted pairwise Kolmogorov-Smirnov tests.

Additionally, we investigated whether the relationship between the possible gain for the partner and participants’ probability to make a prosocial choice can be described in terms of a psychometric function. For the estimation of the psychometric functions we used the R-package “quickpsy” which implements a Maximum-Likelihood-Estimation procedure to fit the cumulative normal distribution. To test whether the points of subjective equality (PSEs) differed between conditions, we conducted a linear mixed model with the condition as fixed effect, participant as random effect and PSE as dependent variable.

To test whether the relative difference between empathy and reciprocity in the *z*-parameter and *a*-parameter could explain the percent changes of these parameters in the multi-motive condition compared to the reciprocity condition, the percent change values (Δ*z*_multi-motive/reciprocity_ and Δ*a*_multi-motive/reciprocity_) were entered as dependent variables in a linear regression model. The respective relative differences (Δ*z*_empathy/reciprocity_ and Δ*a*_empathy/reciprocity_) and one regressor modeling the parameter type (*z*-parameter, *a*-parameter) were included as predictors.

#### 2.6.2. Drift-diffusion modeling

We used hierarchical drift-diffusion modeling (HDDM) (Vandekerckhove et al., 2011; Wiecki et al., 2013), which is a version of the classical drift-diffusion model that exploits between-subject and within-subject variability using Bayesian parameter estimation methods, because it is ideal for use with relatively small sample sizes. The analyses were conducted using the python implementation of HDDM version 0.8.0 (Wiecki et al., 2013).

Based on binary choices, the HDDM approach provides detailed insights into the computation of egoistic and prosocial choices, because it uses all the raw data that is available (trial-by-trial reaction times and choice outcome information of all choices, irrespective of point distributions) to estimate sub-components of the underlying decision process. The v, z and a-parameters for each participant capture how each person maneuvers between the egoistic and the prosocial choice options, and finally approaches a decision boundary (i.e., the boundary for an egoistic or a prosocial choice). In line with previous studies that have used a similar procedure in the realm of social decision making (e.g., Chen and Krajbich, 2018; Gallotti and Grujić, 2019), we believe that the HDDM results provide a sensitive and fine-grained proxy for individual differences in prosociality.

Since we did not have prior hypotheses about which and how many of the three central DDM parameters may reflect motive complexity, we estimated 11 possible variants of the DDM model, ranging from the most simple model (no parameter is modulated by condition) to the full model with *v*, *z*, and *a* possibly being modulated by our four conditions (baseline, empathy, reciprocity, and multi-motive). Since the point information varied between trials, which may influence drift-rate, we allowed the drift rate to vary by the trial-by-trial possible gain for the partner (see section 2.6.1. for computation of this value). We performed model comparison based on the deviance information criterion (DIC) and extracted the parameters of the winning model (lowest DIC value). Apart from the three parameters of interest for our research question (*v, z, a*), additional parameters are included in the estimation procedure. We also estimated the non-decision time t and allowed for trial-by-trial variations of the initial bias (sz), he drift rate (sv) and the non-decision time (st). These parameters were not estimated to vary by condition. They were nonetheless included based on the results by Lerche and Voss (2016), who showed that in most cases, it is beneficial to include these parameters in order to improve model fit. In the estimation procedures we used the default values for the priors and hyperpriors provided by the HDDM package. In more detail, the “informative group mean priors are created to roughly match parameter values reported in the literature and collected by Matzke and Wagenmakers (2009).” cited from (Wiecki et al., 2013, Supplementary Material, page 1). Model convergence was checked by visual inspection of the estimation chain of the posteriors, as well as computing the Gelman-Rubin Geweke statistic for convergence (all values < 1.01) (Gelman and Rubin, 1992). To assess model fit, we conducted posterior predictive checks by comparing the observed data with 500 datasets simulated by our model (Wiecki et al., 2013). This approach allows for the computation of intervals within which the parameter falls with 95 % probability. If the observed data falls within the 95 % credibility interval of the simulated data, the model can describe the data well. The present results revealed a good match between the observed data and the modeled data. Parameters of interest from the winning model were extracted for further analysis. Specifically, for each participant, the condition-specific *v-*parameters, *z-*parameters, and *a-*parameters were extracted (resulting in 12 parameters per participant). In HDDM, the *z*-parameter is always relative to *a*. The reported values of *z* thus range between 0 and 1 and correspond to the absolute value of *z* divided by the a-parameter (*z/a*)

For closer investigation of processing differences in complex vs. more simple motivational states, we compared the posterior distributions of the conditions for each parameter by computing the probabilities for the multi-motive parameter being larger than the single motive parameters. This was done by calculating the densities of the differences distributions that are larger than zero (Wiecki et al., 2013). Additionally, we used the plausible value approach to estimate the corresponding t-value. This approach consists of repeatedly sampling participants’ individual parameters from the winning model’s posterior distribution. Extracting these parameters and comparing between the different conditions using frequentist statistics results in distributions of t-values whose means are a plausible proxy for the actual underlying t-value (Ly et al., 2017; Marsman et al., 2016).

#### 2.6.3. fMRI data acquisition

Imaging data was collected at a 3T MRI-scanner (Verio, Siemens, Erlangen, Germany) with a 32-channel head coil. Functional imaging was performed with a multiband EPI sequence of 72 transversal slices oriented along the subjects’ AC-PC plane (multi-band acceleration factor of 6). The in plane resolution was 2.5 x 2.5 mm² and the slice thickness was 2.5 mm. The field of view was 210 x 210 mm², corresponding to an acquisition matrix of 84 x 84. The repetition time was 1 s, the echo time was 33.6 ms, and the flip angle was 54°. Structural imaging was conducted using a sagittal T1-weighted 3D MPRAGE with 176 slices, and a spatial resolution of 1 x 1 x 1 mm³. The field of view was 250 x 250 mm², corresponding to an acquisition matrix of 256 x 256. The repetition time was 1,690 ms, the echo time was 2.52 ms, the total acquisition time was 3:50 min, and the flip angle was 9°. For the T1-weighted images, GRAPPA with a PAT factor of 2 was used. We obtained, on average, 1,911 (SD = 5.6 volumes) EPI-volumes during the choice task of each participant. We used a rubber foam head restraint to avoid head movements.

#### 2.6.4. fMRI Preprocessing

Preprocessing and statistical parametric mapping were performed with SPM12 (Wellcome Department of Neuroscience, London, UK) and Matlab version 9.2 (MathWorks Inc; Natick, MA). Spatial preprocessing included realignment to the first scan, and unwarping and coregistration to the T1 anatomical volume images. Unwarping of geometrically distorted EPIs was performed using the FieldMap Toolbox. T1-weighted images were segmented to localize grey and white matter, and cerebro-spinal fluid. This segmentation was the basis for the creation of a DARTEL Template and spatial normalization to Montreal Neurological Institute (MNI) space, including smoothing with a 6 mm (full width at half maximum) Gaussian Kernel filter to improve the signal-to-noise-ratio. To correct for low-frequency components, a high-pass filter with a cut-off of 128 s was used.

#### 2.6.5. fMRI statistical analysis

Our participants made prosocial choices in the majority of the trials (Mean = 74 %, SD = 19 %) with more than half of the participants making prosocial choices in 80 % or more of the trials in at least one of the four conditions (see **Table A2**). Given the lack of egoistic choices and given that our study focused on the computation of prosocial choices, egoistic choices trials were not included in the imaging analyses and we also refrained from computing direct contrasts between prosocial and egoistic choices.

First-level analyses were performed with the general linear model (GLM), using a canonical hemodynamic response function (HRF) and its first derivative (time derivative). Regressors were defined from cue onset until the individual response was made by pressing a button (resulting in a time window of 1,000 ms + individual response time). For each of the four conditions (the three motive conditions and baseline condition), the respective regressors of prosocial choice trials were included as regressors of interest. The respective regressors of all the other trials (e.g., egoistic choice trials and trials with missed button presses) were included as regressors of no interest. Given that more than half of our participants (64 %) made fewer than five egoistic decisions in at least one of the conditions, we refrained from computing direct contrasts between prosocial and egoistic choices and included egoistic choices in this regressor of no interest (see **Table A2** for the number of trials per participant and condition). The residual effects of head motions were corrected for by including the six estimated motion parameters for each participant and each session as regressors of no interest. To allow for modeling all the conditions in one GLM, an additional regressor of no interest was included, which modeled the potential effects of session.

For the second-level analyses, contrast images for comparisons of interest (empathy > reciprocity, multi-motive > empathy, reciprocity > empathy, and multi-motive > reciprocity) were initially computed on a single-subject level. In the next step, the individual images of the main contrast of interest (multi-motive > reciprocity) were regressed against the percent change in the *z*-parameter (Δ*z*_multi-motive/reciprocity_) and *a*-parameter (Δ*a*_multi-motive/reciprocity_) in the multi-motive condition, relative to the reciprocity condition, using second-level regressions. Second-level results were corrected for multiple comparisons, using cluster-level family wise error (FWE) correction on a whole brain level. We also report results at a threshold of P_uncorrected_ < 0.001 and a cluster threshold of k > 10 in Appendix A: Supplementary Material.

To test if the neural response in the dorsal striatum was related to the relative difference in *z* between empathy and reciprocity (Δ*z*_empathy/reciprocity_), the (multi-motive > reciprocity) contrast was regressed against the empathy vs reciprocity *z*-differences (Δ*z*_empathy/reciprocity_) and the multi-motive z-enhancement (Δ*z*_multi-motive/reciprocity_) in the same model. Additionally, the individual beta-estimates of the neural multi-motive condition > reciprocity and empathy > reciprocity contrasts were extracted from an independent anatomical ROI of bilateral putamen based on the aal nomenclature (Tzourio-Mazoyer et al., 2002), using MarsBaR (Brett et al., 2002) and the WFU PickAtlas Tool (Maldjian et al., 2003).

In order to clarify the commonly shared influence of the partner’s possible gain on the neural prosocial choice process, we added the partner’s gain as trial-by-trial parametric modulator of the decision phase to a first level GLM in which all conditions are collapsed into one single regressor. On the second level, we conducted a one sample t-test on this parametric modulator corrected for multiple comparisons, using cluster-level family wise error (FWE) correction on a whole brain level.

The reported anatomical regions were identified using the SPM anatomy toolbox (Eickhoff et al., 2005).

### 2.7. Data and code availability

Behavioral data and scripts are available at github.com (https://github.com/AnneSaulin/complex_motivations).

Imaging data are available at neurovault.org (https://www.neurovault.org/collections/5879/).

## 3. Results

### 3.1. Motive induction

During the empathy induction, participants indicated how they felt after observing the person in pain. During the reciprocity induction, they indicated how they felt after receiving a favor from the other person. In the multi-motive condition, participants provided both of these ratings. Strong empathy is indicated by negative feelings when seeing the partner in pain, indicated by negative ratings. Strong reciprocity is indicated by positive feelings when observing the decision of the partner, indicated by positive ratings. To allow the comparison of the ratings in all conditions, empathy ratings were recoded such that positive ratings now reflect strong empathy, i.e., multiplied by -1. The results of linear mixed models (lmms) showed that the induction ratings in the motive conditions were significantly higher than those in the baseline condition (χ^2^ = 515.15, P < .000001, β = 1.61, SE = 0.071, rating_baseline_= -1.02 ± 1.00, rating_empathy_= 1.57 ± 0.77, rating_reciprocity_= 1.50 ± 0.89, rating_multi-motive_ = 1.54 ± 0.91, (*M* ± *SEM*)). There were no significant differences in the induction ratings between the motive conditions (χ^2^ = 0.14, P = .93, β_reciprocity_ = -0.07, SE = 0.20, β_multi-motive_ = -0.02, SE = 0.17). The induction ratings in the motive conditions were significantly associated with the frequency of prosocial choices (χ^2^ = 6.38, P = .01). This effect held to a comparable extent across all three motive conditions (motive condition × rating interaction, χ^2^ = 3.61, P = .16, see **Table A3** for full results). Specifically, the two single-motive conditions yielded similar induction ratings (χ^2^ = 0.23, P = .64, = -0.07, SE = 0.15) and had a comparable effect on the frequency of prosocial choices (χ^2^ = 4.77, P = .03, condition × rating interaction, χ^2^ = 2.06, P = .15, see **Table A4** for full results). These results show that the strength of motive induction and the link to prosocial choices was comparable for the empathy and the reciprocity motives (**Fig. A1**). Further, supporting that the induction procedure specifically influenced empathy and reciprocity motivations, trait empathy and trait reciprocity differentially influenced the frequency of prosocial behavior in the three motive conditions (trait measure type × trait measure value × motive condition interaction, χ = 6.08, P = .047, see **Table A5** for full results).

### 3.2. Frequency of prosocial choices

Pairwise Kolmogorov-Smirnov tests revealed that the frequency of prosocial decisions was comparably distributed across conditions (all Ds < 0.24, all Ps > 0.29, for detailed statistics, please see **Table A6**) as were the reaction times (all Ds < 0.27, all Ps > .17, **Fig. 3C**).

**Figure 3.**
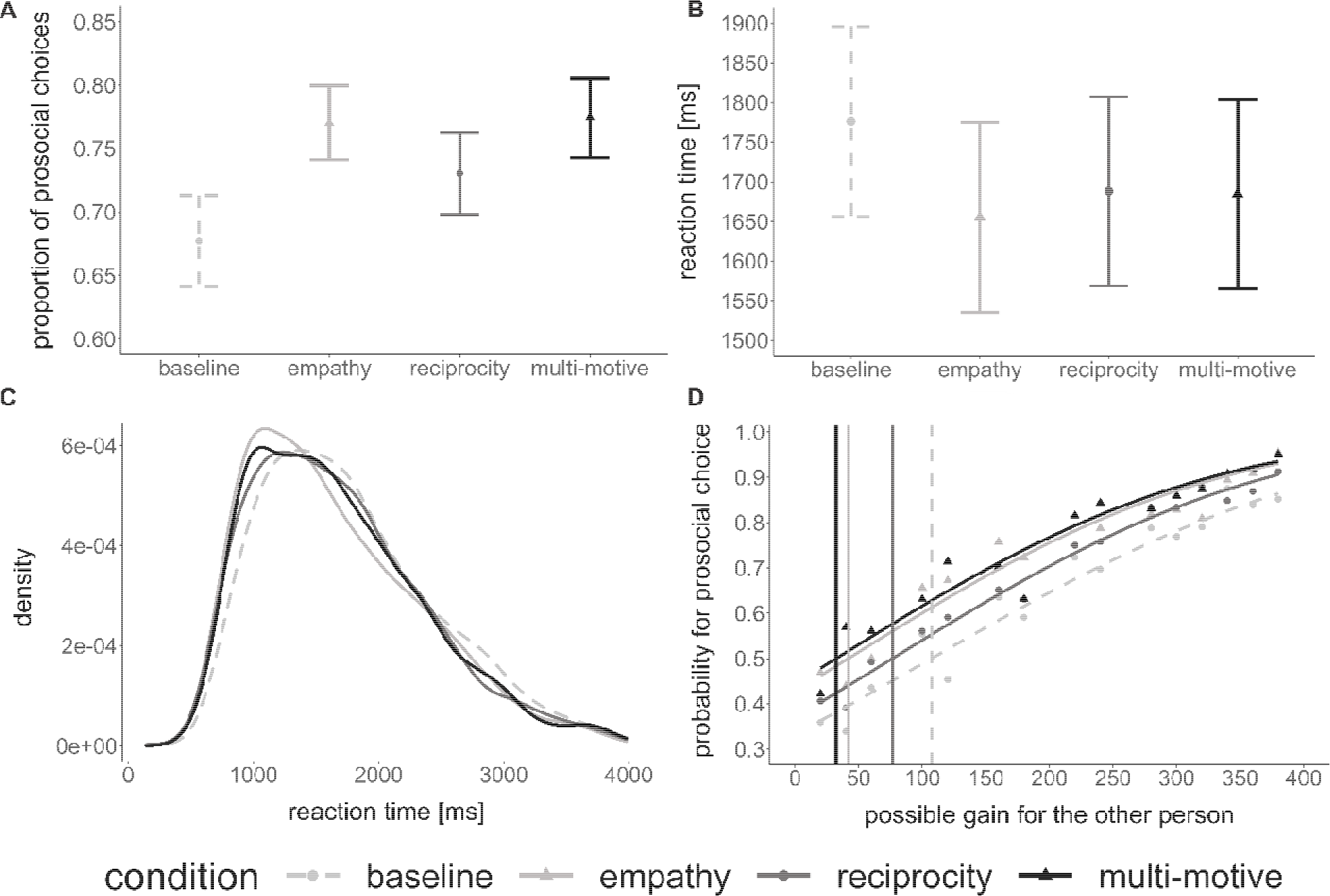
Descriptive statistics, distributions and psychometric function of the choice and reaction time data. **(A)** Mean proportion of prosocial choices per condition. Error bars denote standard errors of means. **(B)** Mean reaction times per condition. Error bars denote standard errors of means. **(C)** Distribution of reaction times across participants per condition. **(D)** Psychometric functions for the different conditions of the probability to make a prosocial choice depending on the amount of points the participants’ partner could possibly gain in each trial, that is, the point value for the partner in case of a prosocial choice minus the point value for the partner in case of an egoistic choice. Vertical dashed lines indicate the points of subjective equality in the different conditions (for exact values and spread, see **Table A9**). Please note that the probability of making a prosocial decision never reached 0 because of the high frequency of prosocial choices in our data (participants made prosocial choices even if the gain for the other person was low). Thus, our results do not yield data points much lower than the respective points of subjective equality (PSEs).

The frequency of prosocial choice was significantly influenced by condition (χ^2^ = 56.99, P < .0001, see **Fig. 3A** and **Fig. A2**, prosoc_baseline_= 67.7 ± 20.6 %, prosoc_empathy_= 77.0 ± 16.8, prosoc_reciprocity_= 73.1 ± 18.4, prosoc_multi-motive_= 77.4 ± 18.0 (*M* ± *SEM*), see **Table A7** for full results), indicating that the motive inductions had a differential effect on later prosocial choices. Moreover, prosocial choices were influenced by the possible gain for the partner (χ^2^ = 668.64, P < .0001). However, model comparison revealed that neither including the possible gain for the participant (χ^2^ = .23, P = .63), nor its interaction with condition significantly improved the model fit (χ^2^ = .86, P = .84). Thus, for the analyses reported below condition and possible gain for the partner were used as additional predictors.

The frequency of empathy-based prosocial choices was increased compared to reciprocity-based choices (z ratio = 2.94, P = .02), whereas the frequency of prosocial choices between the multi-motive condition and the empathy condition was comparable (z-ratio = .56, P = .94). However, the multi-motive condition yielded significantly more prosocial choices compared to the reciprocity condition (z-ratio = 3.49, P = .003).

To clarify this effect, we calculated the percent change in prosocial choices in the multi-motive condition relative to each single motive condition

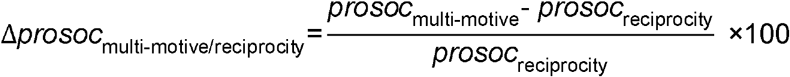

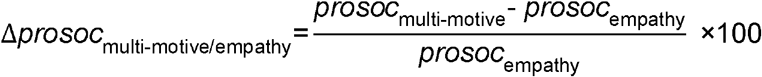

where *prosoc*_multi-motive_ equals the frequency of the prosocial choices in the multi-motive condition, *prosoc*_reciprocity_ equals the frequency of prosocial choices in the reciprocity condition, and *prosoc*_empathy_ equals the frequency of prosocial choices in the empathy condition.

The percent change of the multi-motive condition relative to reciprocity was significantly positive (*t*(32) = 2.07, P = .047, Δ*prosoc*_multive/reciprocity_ = 8.61 ± 4.17 (*M* ± *SEM*)), demonstrating that prosocial choices were enhanced when reciprocity was combined with empathy, relative to reciprocity alone. The percent change in the multi-motive condition relative to the empathy condition was not significantly different from zero (*t*(32) = 0.42, P = .674, Δ*prosoc*_multi-motive/empathy_ = 1.05 ± 2.47 (*M* ± *SEM*)), multi-motive/empathy indicating that the simultaneous activation of the reciprocity motive did not enhance the empathy motive.

### 3.3. Reaction times

Reaction times were significantly influenced by conditions (χ^2^ = 27.89, P < .0001, see **Table A8** for full results). That is, participants were faster in the motive conditions compared to the baseline condition (baseline vs. empathy: t(32) = 5.03, P < .0001, baseline vs. reciprocity: t(32) = 3.62, P = .002, baseline vs. multi-motive: t(32) = 3.70, P = .001). There were no differences in reaction times for prosocial choices between the motive conditions (all Ps > .49) (**Fig. 3B**), and the reaction time distributions were comparable (**Fig. 3C**).

### 3.4. Point equality and prosocial behavior

In a next step, we tested whether considerations of equity differentially influenced participants’ prosocial behavior in the different conditions.

We considered the difference in point equality of the participant’s and the partner’s outcome between the two choice options to test whether inequity aversion differentially influenced participants’ prosocial choice behavior. The results showed a main effect of difference in point equality (χ^2^ = 65.87, P < .0001) and a main effect of condition (χ^2^ = 46.91, P < .0001). However, no interaction effect was observed (χ^2^ = 0.19, P = .98, see **Table A10** for full results). Based on these results we conclude that inequity aversion does not differentially affect the different motive conditions and thus cannot explain behavioral differences in the different conditions.

In line with these results, the relationship between the frequency of prosocial choices and the partner’s possible gain was comparable between the different motive conditions, as reflected by comparable values for the points of subjective equality based on the psychometric functions estimated for the different conditions (χ^2^ = 2.89, P = .41, **Fig. 3D** and **Table A9**).

Since we were most interested in the underlying prosocial decision processes in more complex as compared to simpler motivational states, we used hierarchical drift-diffusion modeling (HDDM) (Vandekerckhove et al., 2011; Wiecki et al., 2013) to understand prosocial choice behavior in the multi-motive condition relative to the reciprocity condition and relative to the empathy condition.

### 3.5. Hierarchical drift-diffusion modeling

We estimated the three aforementioned DDM parameters (*v, z, a*) for every condition and participant. We also estimated the non-decision time *t* (0.94 ± 0.04 (*M* ± *SEM*)). However, this parameter was not estimated to vary by condition and was thus not further analyzed. Comparing the observed data with 500 datasets simulated by our model (Wiecki et al., 2013) showed that the winning model fit the data with 95% credibility (see **Table A11** for overview of all models and DIC values, and see **Table A12** for quantile comparison and 95% credibility). Based on the hypotheses depicted in **Fig. 1C**, we tested whether the observed percent change in the multi-motive condition can be explained by an increase in the speed of information accumulation (*v*-parameter, **Fig. 1C**, left panel), and/or an increase in initial prosocial bias (*z*-parameter, **Fig. 1C**, middle panel). Additionally, we tested whether the induction of both motives enhanced the amount of relative evidence that participants required during the choice process, relative to the two single-motive conditions (*a*-parameter, **Fig. 1C**, right panel).

Testing the first hypothesis (**Fig. 1C**, left panel), we observed no significant differences between the motive conditions (*p(v_multi-motive_>v_empathy_) = 46.38 %,* plausible t(32) = -.31, P = .76; *p(v_multi-motive_>v_reciprocity_) = 72.25 %,* plausible t(32) = 1.03, P = .31, see **Fig. A4** for distribution of t-values). Further, there was a slight percent change in *v*-parameters in the multi-motive condition relative to the reciprocity condition
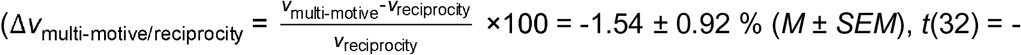
1.71, P = .09) and no percent change relative to the empathy condition
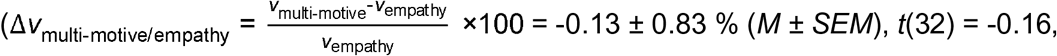
P = .88). This result showed that the speed of information accumulation, i.e., the efficiency of the choice process itself, was mainly unaffected by the combination of the two motives, relative to the single-motive conditions.

Testing the second hypothesis (**Fig. 1C**, middle panel), we observed an increase in initial prosocial bias (*z*-parameter) in the multi-motive condition compared to the reciprocity condition (*p(z_multi-motive_>z_reciprocity_) = 93.55 %,* plausible t(32) = 3.66, P < .001) (**Fig. 4A**), but not compared to the empathy condition (*p(z_multi-motive_>z_empathy_) = 70.33 %*, plausible t(32) = 1.67, P = .10). The percent change in the *z*-parameter of the multi-motive condition was significantly positive relative to the reciprocity
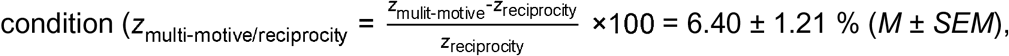 (*t*(32) = 5.36, P < .001) and marginally larger than zero relative to the empathy
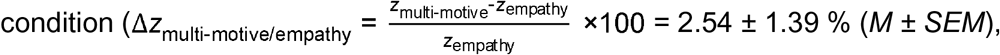 (*t*(32) = 1.85, P = .07).

**Figure 4.**
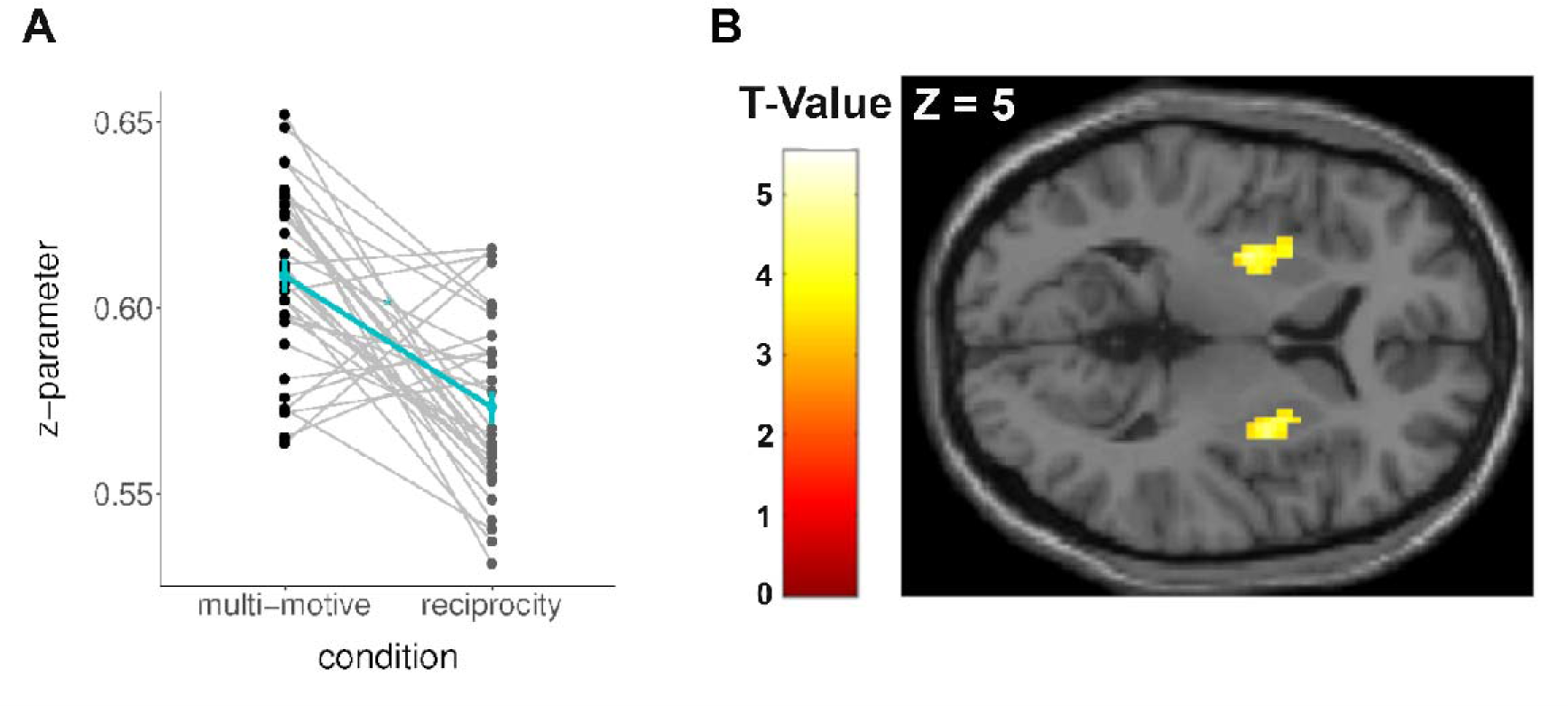
Increase in initial prosocial bias in the multi-motive condition relative to the reciprocity condition and related neural activity. **(A)** Initial prosocial bias (*z*-parameter) were significantly stronger in the multi-motive compared to the reciprocity condition (plausible t(32) = 3.66, P < .001). Individual values are depicted for the multi-motive condition (red) and the reciprocity condition (blue). Means and standard errors of the mean are depicted in black. **(B)** The individual changes of initial prosocial choice biases in the multi-motive condition relative to the reciprocity condition were tracked by an increase in neural responses in the bilateral dorsal striatum (P(whole-brain FWE_cluster-corrected_) = .001; MNI peak coordinates; right hemisphere: x = 30, y = 2, z = - 2, left hemisphere: x = -28, y = -9, z = 1; visualized at P < .001 uncorrected; **Table A14** and **Fig. A6**).

In addition, we had hypothesized that the combination of the two motives may increase the amount of relative evidence that participants required in order to reach a decision (captured by the *a*-parameter; **Fig. 1C**, right panel). The *a*-parameter was not significantly higher in the multi-motive condition compared to the reciprocity condition (*p(a_multi-motive_>a_reciprocity_) =* 84.70 %, plausible t(32) = 1.73, P = .09) and the empathy condition (*p(a_multi-motive_>a_empathy_) =* 82.35 %, plausible t(32) = 1.43, P = .16). However, there was a significantly positive relative percent change in *a*-parameters in the multi-motive condition relative to the reciprocity condition
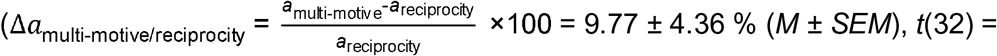
2.28, P = .03) and also relative to the empathy condition
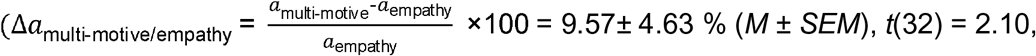 P = .04).

Taken together, the DDM results showed that the combination of the two motives enhanced participants’ bias for choosing the prosocial option, relative to the initial prosocial choice bias biases induced by the reciprocity motive (captured by the percent change in the *z*-parameter). The combination of empathy and reciprocity also led to a relative increase in the amount of relative evidence that people required to make a choice relative to the reciprocity motive, and also relative to empathy (captured by the percent change in the *a*-parameter). In contrast, the speed of information accumulation, i.e., the efficiency of the choice process itself, was comparable between multi-motive and single-motive conditions (no change in *v*-parameter).

These results may indicate that the observed percent changes in the multi-motive condition relative to the reciprocity condition (in the *z-* and the *a-*parameters) originate from the simultaneous activation of the two motives in the multi-motive condition. Alternatively, as we observed no significant difference between the multi-motive condition and the empathy-condition, it is also conceivable that the empathy motive replaced the reciprocity motive when the two motives were activated simultaneously. In this case, the observed percent changes in the multi-motive condition would reflect the dominance of empathy over reciprocity, instead of a multi-motive effect. If in fact empathy replaced the co-activated reciprocity motive, the relative difference in the *z*-parameters and *a*-parameters between the empathy and the reciprocity conditions should predict the individual extent of the percent changes in the multi-motive condition relative to the reciprocity condition. To test this explanation, we calculated the relative differences in the *z*-parameters and *a-* parameters between empathy and reciprocity
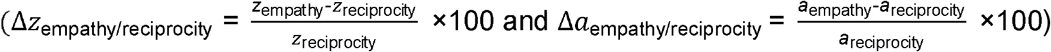, entered them as predictors in a regression analysis, and tested their effects on the observed percent changes in the multi-motive condition (*z*_multi-motive/reciprocity_; Δ*a*_multi-motive/reciprocity_). This analysis revealed no significant effects (β = -0.20, SE = 0.255, P = .42; interaction with parameter type (*z* vs *a*): β = .002, SE = 0.38, P = 1.00, main effect of empathy dominance: β = 0.08, SE = 0.136, P = .59). These results demonstrate that the difference between the two single motives cannot account for the changes in choice parameters in the multi-motive condition relative to the reciprocity condition, bolstering the claim that the observed effects are driven by the simultaneous activation of the two motives. The three DDM parameters of interest for each condition and the relative differences between the baseline condition and the motive conditions are provided in **Figure A3** and **Table A13**.

### 3.6. Imaging results

Next, we investigated the neural underpinnings of the prosocial decision process comparing the multi-motive and the single motive conditions. The main contrasts of mean neural activation during the prosocial decision phase did not show significant neural activations (neither whole-brain nor small-volume corrected).

We investigated how the simultaneous activation of the two motives, and the resulting changes in initial prosocial bias and amount of required relative evidence affected the neural computation of prosocial choices. To do so, we regressed participants’ individual percent change in initial prosocial biases (Δ*z*_multi-motive/reciprocity_) and the amount of relative evidence (Δ*a*_multi-motive/reciprocity_) on the neural contrast in prosocial choices between the multi-motive condition and the reciprocity condition, using second-level regressions. As a main result, the first analysis revealed activations in the bilateral dorsal striatum that were related to the individual change in prosocial bias (right hemisphere: P(FWE_cluster-corrected_) = 0.001; center co-ordinates: x = 30, y = 2, z = -2; k = 143 voxels, t(31) = 5.49; left hemisphere: P(FWE_cluster-corrected_) = 0.003; center co-ordinates: x = -28, y = -9, z = 1; k = 121 voxels, t(31) = 5.36; **Fig. 4B, Fig. A6, Table A14).** The stronger the percent increase in initial prosocial bias in the multi-motive condition relative the reciprocity condition, the stronger the neural response in bilateral dorsal striatum.

To test the alternative hypothesis that the increase in dorso-striatal activity may reflect the dominance of empathy (captured by the relative difference in *z*-parameters between empathy and reciprocity, Δ*z*_empathy/reciprocity_), instead of a multi-motive effect, we also compared the relationship between Δ*z*_multi-motive/reciprocity_ and Δ*z*_empathy/reciprocity_ on extracted beta values of the multi-motive vs reciprocity contrast using an independent anatomical mask of bilateral putamen based on the aal nomenclature (Tzourio-Mazoyer et al., 2002). The results showed that neural activation in dorsal striatum is associated with Δ*z*_multi-motive/reciprocity_, but not with Δ*z*_empathy/reciprocity_ (significant interaction between index type and neural activation: β = 0.69, SE = 0.225, P = .003, no main effect of beta values: β = -0.11, SE = 0.159, P = .52, marginal main effect of index type: β = -0.41, SE = 0.223, P = .07, **Fig. A5B**). Thus, empathy dominance is not likely to explain the results.

To test whether the increase in striatal activation in the multi-motive compared to the reciprocity condition is driven by outliers, we extracted the individual beta-estimates of the multi-motive vs reciprocity contrast from bilateral dorsal striatum and plotted its relationship with the percent change in the z-parameter in the multi-motive condition relative to the reciprocity condition (**Fig. A5A**). The inspection of the plot shows that the relationship between the percent signal change in the z-parameter in the multi-motive condition relative to the reciprocity condition was not driven by outliers.

The respective second-level regression with the percent change in the *a*-parameter revealed neural activity in bilateral anterior insula on a lower, uncorrected threshold (P_uncorrected_ < 0.001; center co-ordinates right hemisphere: x = 33, y = 32, z = 1, P(FWE_cluster-corrected_) = .970, k = 9 voxels; center co-ordinates left hemisphere: x = -30, y = 27, z = -2, P(FWE_cluster-corrected_) = .902, k = 13 voxels).

Additionally, we tested whether trial-by-trial changes in the partner’s gain modulate neural activation during the prosocial choice process. In line with the behavioral results, the partner’s gain did not differentially influence neural activation in the different conditions. However, neural activation during the prosocial choice process in bilateral insula was significantly associated with trial-by-trial changes in the partner’s gain across all four conditions (right insula peak-coordinates: x = 43, y = -6, z = 18, k = 108 voxels, P(FWE whole-brain cluster corrected) = .009; left insula peak-coordinates: x = -38, y= -9, z = 16, k = 517 voxels, P(FWE whole-brain cluster corrected) < .001; see **Table A15** for all clusters k > 10). The same analysis including prosocial as well as egoistic choice trials replicated this result (right insula peak-coordinates: x = 43, y = -6, z = 16, k = 109 voxels, P(FWE whole-brain cluster corrected) = .009; left insula peak-coordinates: x = -38, y= -9, z = 18, k = 294 voxels, P(FWE whole-brain cluster corrected) < .001). Hence, the insular activation appears to track trial-by-trial changes of the partner’s gain across all conditions.

## 4. Discussion

Many behaviors derive from complex motivational states that are characterized by different, simultaneously activated motives (Engel and Zhurakhovska, 2016; Hughes and Zaki, 2015; Jagers et al., 2017; Takeuchi et al., 2015; Terlecki and Buckner, 2015). However, the mechanisms through which combinations of motives affect behaviors, e.g., the computation of prosocial choices, are poorly understood.

Our results showed that the simultaneous activation of two motives changes participants’ choices compared to activation of a single motive condition. In more detail, a combination of two prosocial motives (the empathy and the reciprocity motive) elicited more prosocial choices than the reciprocity motive alone (**Fig. 3A**). This multi-motive increase occurred although the two single motives were activated with comparable strength (indicated by the induction ratings, **Fig. A1**). Moreover, the different motive conditions had no effect on reaction times (**Fig. 3B** and **3C**), inequality aversion, or the subjective value assigned to the partner’s gains (**Fig. 3D**). Furthermore, the partner’s gain was associated with neural activation in bilateral insula across all conditions, which adds to the observation that insular activation is also sensitive to other-regarding experiences such as vicarious reward (Morelli et al., 2015), avoiding risk for others (Shenhav and Greene, 2010), and making fair (Dawes et al., 2012) or altruistic decisions (Cutler and Campbell-Meiklejohn, 2019).

Specifying the change in prosocial choice behavior in the multi-motive condition, the drift-diffusion-modeling analyses showed that the combination of the two motives enhanced participants’ initial bias for making a prosocial choice, compared to the reciprocity condition (reflected by the increase in *z*-parameter; **Fig. 4A**) and with a similar trend, relative to the empathy condition. Moreover, the combinations of the two motives increased the relative amount of relative evidence that participants required during the choice process (reflected by the relative increase of the *a*-parameter). This indicates that participants assessed their choices more carefully if they made them based on two different motives. In contrast, the speed of information accumulation, i.e., the efficiency of the decision process itself (reflected by the *v*-parameter), remained unchanged. The observed change in initial prosocial bias (the *z*-parameter) is in line with previous findings that reported a shift of choice biases due to the prior likelihood of one of the choice options or a higher reward value associated with one option (Mulder et al., 2012), personal predispositions (Chen and Krajbich, 2018), or prior information about how other people decided (Toelch et al., 2018). Extending these results, our findings reveal that initial choice biases are altered by simultaneously activated motives, and thus characterize how complex motivational states change the choice process compared to single-motive states.

We hypothesized that changes in DDM choice parameters in the multi-motive compared to the single-motive conditions might be related to changes in activation in the ventral or dorsal striatum, inspired by evidence associating the ventral striatum with the processing of choice values (Kable and Glimcher, 2007; Liljeholm and O ’Doherty, 2012; O’Doherty et al., 2004; Strait et al., 2015), and/or the dorsal striatum with encoding of choice preferences (Balleine et al., 2007; Liljeholm and O ’Doherty, 2012; O’Doherty et al., 2004; Palmiter, 2008; Robinson et al., 2006).Our results show that the combination of different motives is associated with an increase in activation in bilateral dorsal striatum, reflecting an enhancement of individual prosocial choice biases in the multi-motive condition relative to the reciprocity condition (**Fig. 4**), and, based on extracted beta-values from an independent anatomical region, also relative to the empathy condition (**Fig. A5**). The increase in activation in dorsal striatum is in line with previous neuroscience studies showing that motivation-related changes in decision parameters are captured by dorsal striatal responses (Forstmann et al., 2008; Gluth et al., 2012). Extending this previous evidence, we show that the dorsal striatum integrates choice biases that are elicited by multiple motivational forces, and thus provides a plausible neural candidate for the generation of complex motivational states.

We found that the simultaneous activation of the empathy motive and the reciprocity motive in the multi-motive condition enhanced the participants’ initial prosocial biases relative to the reciprocity condition. This indicates that the empathy motive enhanced the reciprocity motive, but not vice versa. Given this result, we argued that the observed changes in the multi-motive condition may reflect the dominance of one motive over the other motive (i.e., a dominance of empathy over reciprocity). If this were true, the multi-motive induced changes in the choice process would reflect a motivation that is similar to the state induced by the dominant motive, instead of a more complex motivational state that was incited by the combination of different motives. Our results show that the multi-motive induced changes in the choice process (i.e., DDM and neural choice parameters) are in fact related to differences between the multi-motive condition and the reciprocity condition and cannot be explained by mere dominance of the empathy motive over the reciprocity motive. This finding supports the conclusion that the simultaneous activation of two motives alters the prosocial choice process compared to single motive states.

Because we were mainly interested in participants’ choices under the different motive conditions, our paradigm was designed to optimize the number of trials in the allocation task. The motive induction procedures only included the minimal number of trials required for inducing the different motives (twelve trials per motive induction). Due to the small number of trials, an analysis of neural responses during the multi-motive and single-motive induction procedures would not be meaningful.

Participants made prosocial choices also in the baseline condition which indicates that participants are motivated to behave prosocially without experimental activation of empathy and reciprocity. It is thus important to note that prosocial choices can be driven by other motives in addition to empathy and reciprocity. However, these additional motives should be the same across conditions since participants perform the same social choice task. Hence, contrasting behavior between the different conditions should carve out the effects that were experimentally manipulated, i.e., how the combination of empathy and reciprocity influences the prosocial choice process relative to empathy or reciprocity alone.

Likewise, because the “basic” social choice tasks were identical in all experimental conditions, task-specific effects should average out if the different conditions are contrasted. This means that the observed effects are driven by the different motive inductions and should be independent of the social choice task that was used in the present study. In other words, our behavioural and neural findings should generalize to other behaviours that are elicited by the combination of the empathy and the reciprocity motives. However, in how far the present effects are scaled depending on the exact task affordances (e.g., relative importance of the single motives for the respective task, how much time participants have to deliberate their decision) is a question for future research. Likewise, future studies need to test if the observed increase in striatal activation due to a multi-motive alteration of initial choice bias also applies to other (e.g., non-social) motivational states. Food choices, for example, are often driven by more than one motive such as the motive to eat healthy and the motive to eat sweet food, maximizing calorie intake. The dorsal striatum has previously been associated with food choice preferences in healthy (Small et al., 2003; Wallace et al., 2014) as well as pathological participants (Foerde et al., 2015). It is thus possible that the interplay between the non-social motives during food choices is associated with neural activation in the dorsal striatum.

To avoid cross-gender effects, which are likely to occur if female participants are paired with male confederates and vice versa, we only tested females. Future studies are required to show if our results generalize to male participants.

## 5. Conclusions

Based on our current findings we conclude that the simultaneous activation of two different prosocial motives changes the computation of prosocial choices. According to our results, choices that are made in a more complex motivational state, i.e., driven by multiple motives, are characterized by a change in initial choice bias, which is associated with an increased neural response in dorsal striatum. Moreover, choices were made more carefully relative to simple motivational states. Together, these findings show how the human brain combines different prosocial motives, and how this motive combination affects the computation of prosocial choices.

## Authors’ Contributions

G.H. and A.S. designed the research with input from J.K., M.L., and U.H.; A.S. and U.H. performed the research; A.S. programmed the experiment; A.S. and U.H. analyzed the data with input from G.H., M.L., and J.K.; G.H. and A.S. wrote the paper with input from U.H., M.L., and J.K..

## Acknowledgments

We thank Martin Domin, Jörg Pfannmöller, and Kai Klepzig for their technical support, and Vassil Iotzov for helpful discussions about the data.

## Funding

This work was supported by the German Research Foundation (HE 4566/2-1; HE 4566/5-1) and the German Academic Scholarship Foundation.

## Notes

https://github.com/AnneSaulin/complex_motivations

https://www.neurovault.org/collections/5879/

### Competing Interest Statement

The authors have declared no competing interest.

### Summary of Updates

More detailed analyses and regression tables have been added

