## SupplementaryMaterial for "The neural computation of human prosocial choices in complex motivational states"

**Appendix A: Supplementary Material**

**Supplementary Figures**

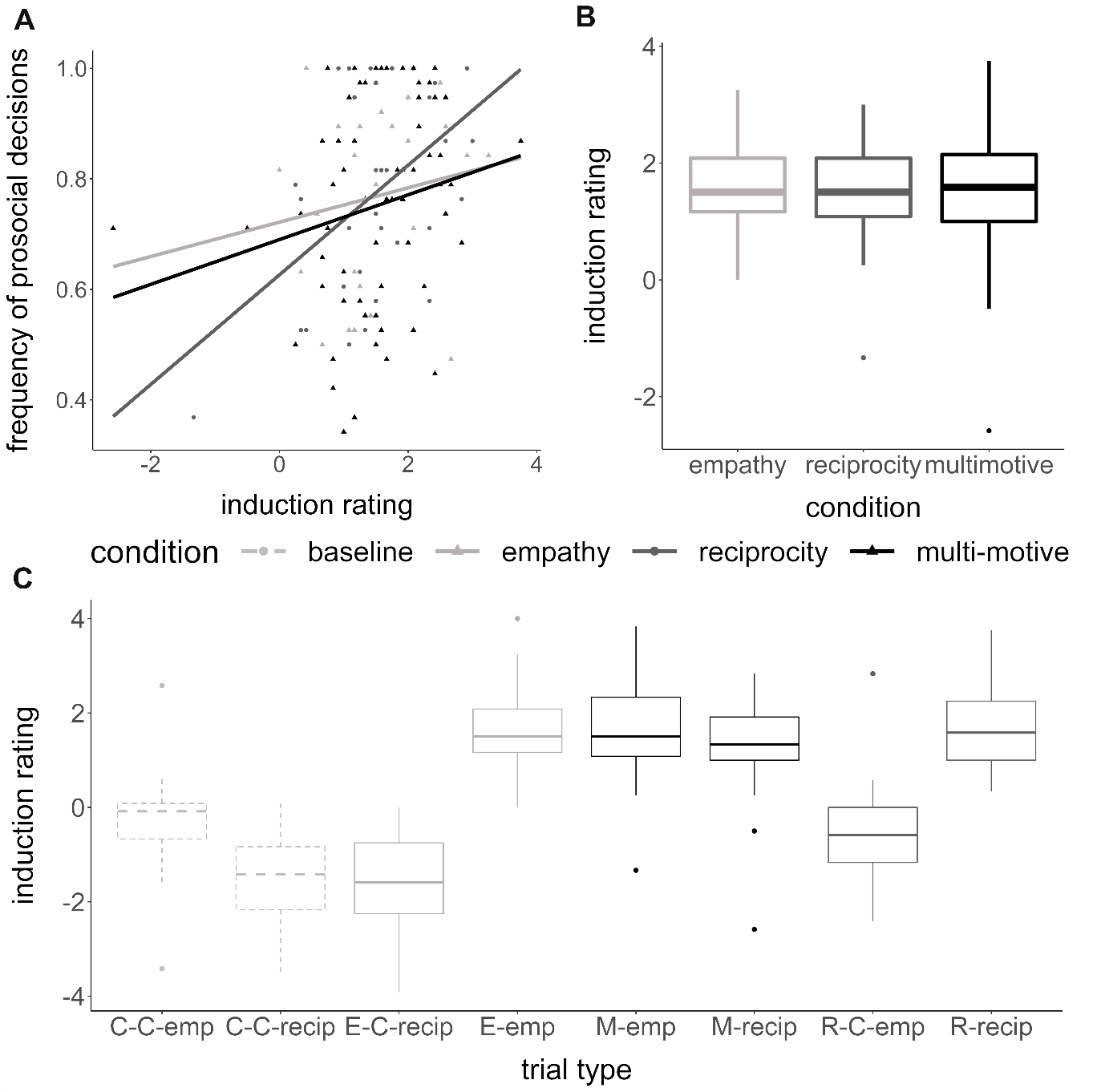

**Figure A1**. Induction ratings in the different motive conditions and trial types and their relationship with the frequency of prosocial choices. (A) Relationship between the mean induction ratings in the motive conditions during motive induction and the respective frequency of prosocial choices in the allocation task. (B) Induction ratings per motive condition. Averages and spread are visualized using boxplots indicating the median and 25th and 75th percentile. (C) Mean induction ratings per trial type of the motive induction. One can see that all trials in which either the other person only receives non-painful stimulations (C-C-emp and R-C-emp) or the computer makes the choices to “help” or not (C-C-recip and E-C-recip) yield lower induction values than the treatment trials with 75% high pain for the other person (E-emp and M-emp) and an actual person making the decision to help or not to help (M-recip and R-recip). C-C-emp/C-C-recip: empathy control trials (only no pain) and reciprocity control trials (computer decides) in the baseline condition; E-C-recip/E-emp: reciprocity control trials (computer makes decision) and empathy trials in the empathy condition; M-emp/M-recip: empathy and reciprocity trials in the multi-motive condition; R-C-emp/R-recip: empathy control trials (only no pain) and reciprocity trials in the reciprocity condition.

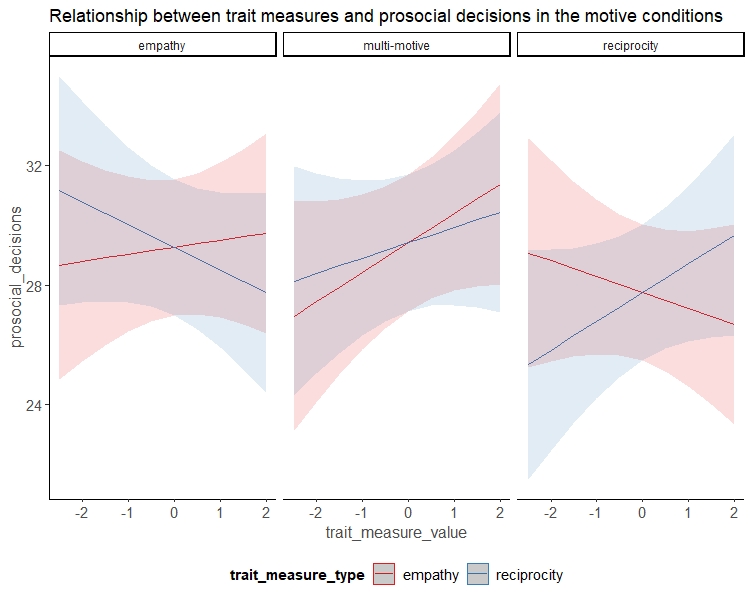

**Figure A2**. The relationship between the number of prosocial choices and the different trait measure types (empathy trait measure vs. reciprocity trait measure) differentially depends on the motive condition (empathy, multi-motive, reciprocity) (three-way interaction of trait measure type, trait measure value, and condition; χ^2^ = 6.08, P = .047).

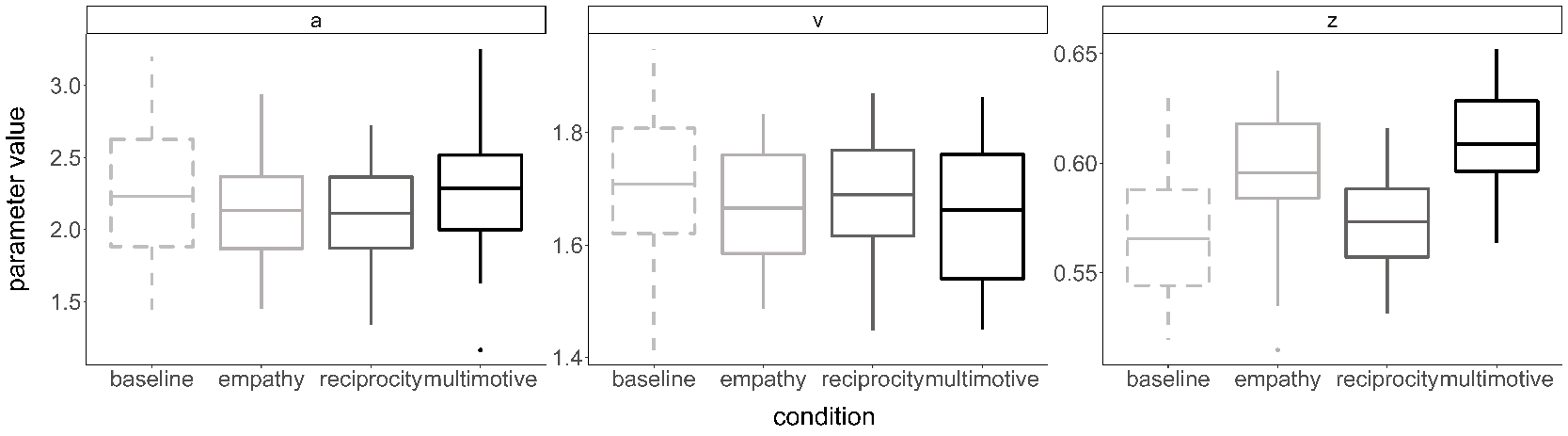
**Figure A3**. Mean values and spread of the parameter estimates for all three model parameters by condition. The *a*-parameter reflects the amount of required relative evidence in order to reach a decision (left panel), the *v*-parameter reflects the speed of evidence accumulation (middle panel), and the *z*-parameter reflects the initial prosocial bias (right panel).

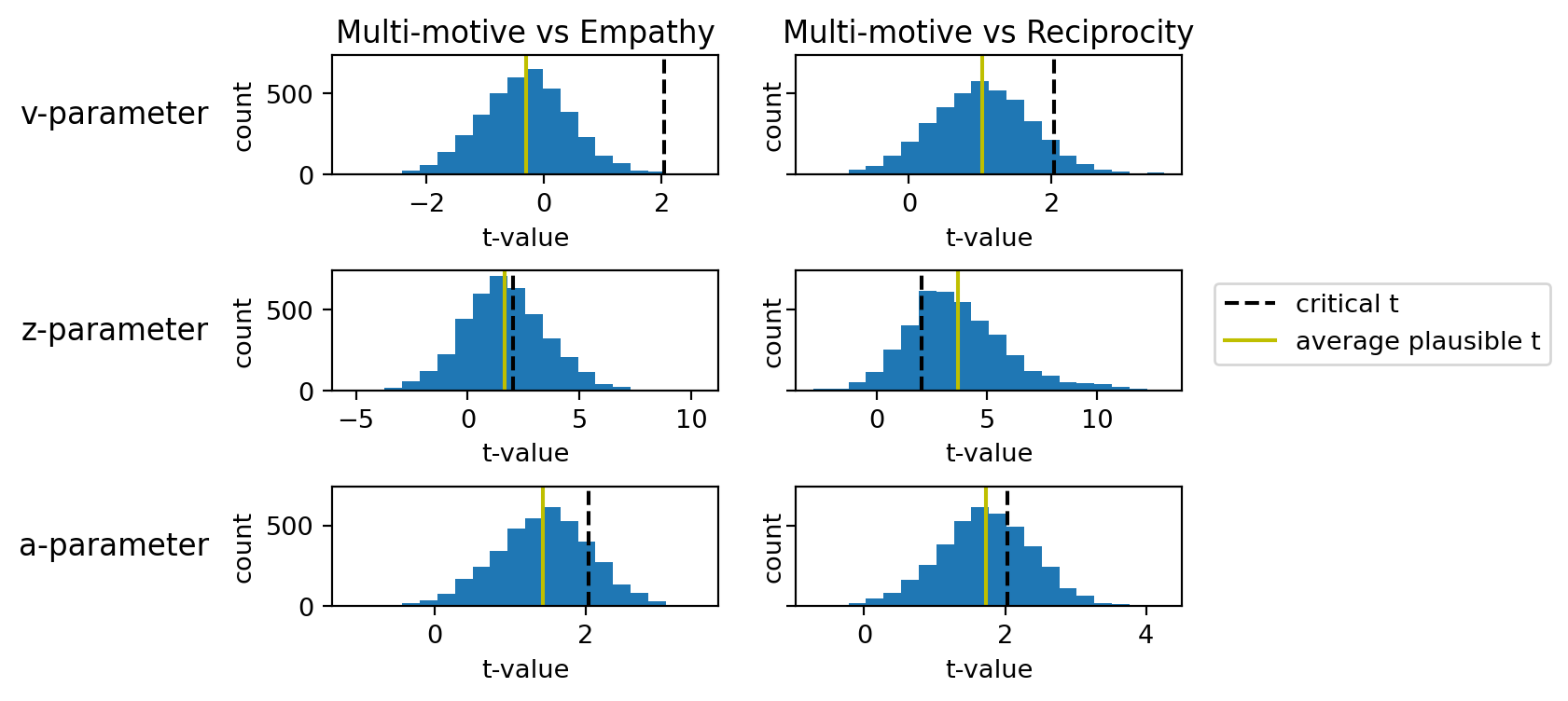
**Figure A4.** Means and distributions of t-values resulting from the plausible values approach comparing the three drift-diffusion model parameters in the multi-motive condition with those in the two single-motive conditions. The black dashed line marks the t-value which corresponds to P < .05 (two-sided) and the green line the respective mean plausible t-value.

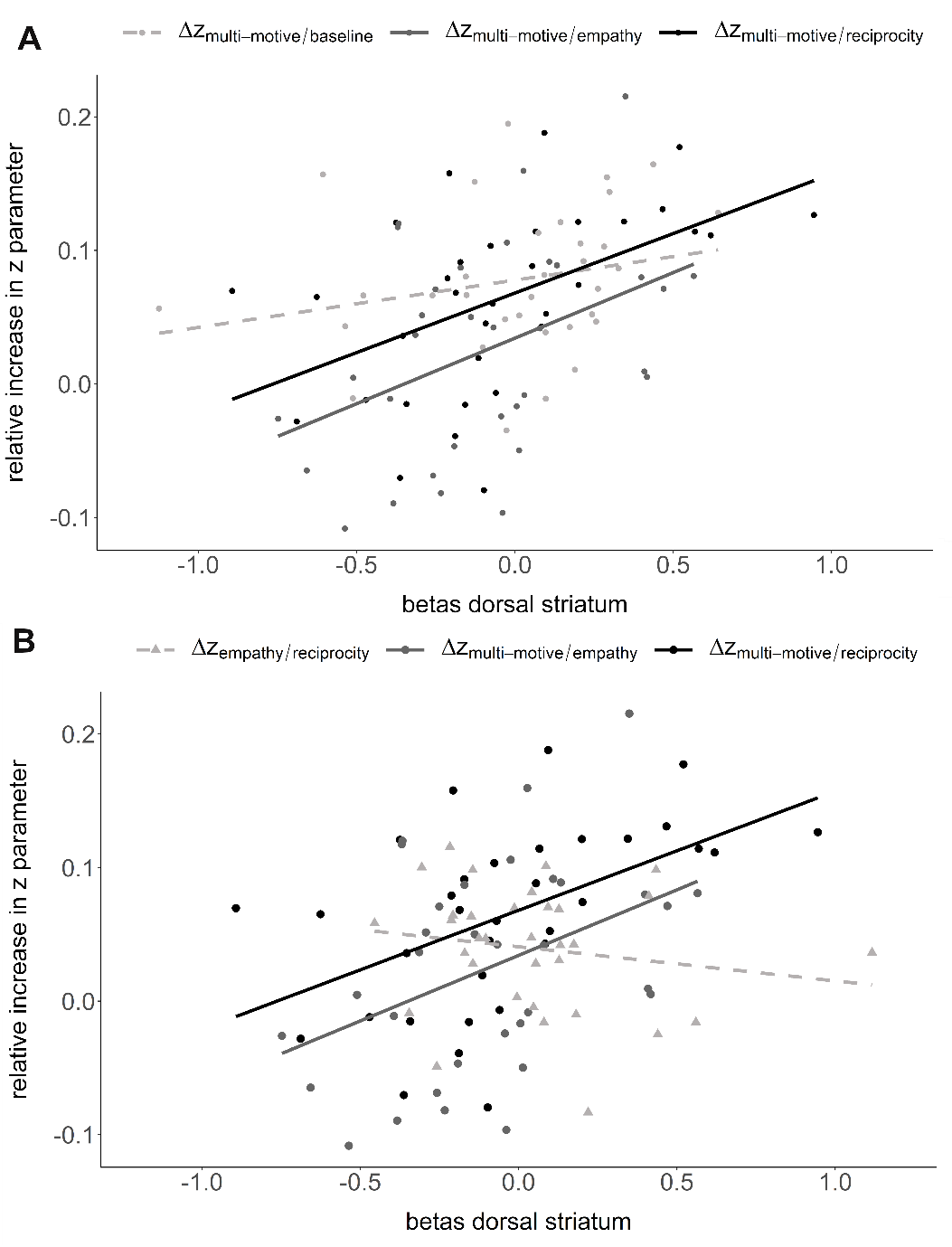

**Figure A5**. Scatterplots visualizing the relationship between the relative increase in z-parameter and the corresponding neural activation in dorsal striatum. **(A)** Scatterplot visualizing the relationship between neural activation in dorsal striatum and the relative increase in the multi-motive *z*-parameter relative to the baseline condition (neural contrast multi-motive > baseline, $\Delta\text{z}_{\text{multi-motive/baseline}}$, lightgrey, rho = .37, P = .03), relative to reciprocity (neural contrast multi-motive > reciprocity, $\Delta\text{z}_{\text{multi-motive/reciprocity}}$, black, rho = .54, P = .001), and relative to empathy (neural contrast multi-motive > empathy, $\Delta\text{z}_{\text{multi-motive/empathy}}$, darkgrey, rho = .41, P = .02). **(B)** Scatterplot visualizing the relationship between neural activation in dorsal striatum and the relative increase in the multi-motive *z*-parameter relative to reciprocity and relative to empathy, as well as the relative increase in the empathy *z*-parameter relative to reciprocity (neural contrast empathy > reciprocity, $\Delta\text{z}_{\text{empathy/reciprocity}}$, lightgrey, rho = -.22, P = .21). The results indicate that empathy dominance is not likely to explain the neural effect in dorsal striatum. Betas were extracted from an independent anatomical mask of bilateral putamen based on the aal nomenclature.

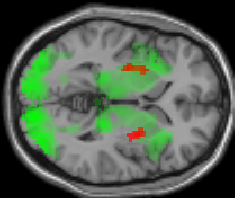
**Figure A6**. Overlap of the neural activations during the decision phase in the multi-motive condition and the reciprocity condition (green) with the neural activation in bilateral putamen based on the second level regression with the relative increase in initial prosocial biases (red). P<.001 uncorr.

**Supplementary Tables**

**Table A1**. Overview of the point options. In both options, the participant’s absolute gain was always larger than the partner’s gain. The prosocial option, however, maximized the partner’s gain at a cost to the participant.

|  | Egoistic option | | Prosocial option | |
| --- | --- | --- | --- | --- |
| distribution | participant gain | partner gain | participant gain | partner gain |
| 1 | 1020 | 0 | 1000 | 380 |
| 2 | 1030 | 0 | 990 | 380 |
| 3 | 1040 | 0 | 980 | 380 |
| 4 | 1060 | 0 | 960 | 380 |
| 5 | 1070 | 10 | 950 | 370 |
| 6 | 1090 | 10 | 930 | 370 |
| 7 | 1100 | 20 | 920 | 360 |
| 8 | 1120 | 30 | 900 | 350 |
| 9 | 1130 | 40 | 890 | 340 |
| 10 | 1150 | 50 | 870 | 330 |
| 11 | 1160 | 70 | 860 | 310 |
| 12 | 1170 | 80 | 850 | 300 |
| 13 | 1180 | 100 | 840 | 280 |
| 14 | 1190 | 110 | 830 | 270 |
| 15 | 1190 | 130 | 830 | 250 |
| 16 | 1200 | 140 | 820 | 240 |
| 17 | 1200 | 160 | 820 | 220 |
| 18 | 1200 | 170 | 820 | 210 |
| 19 | 1200 | 180 | 820 | 200 |
| 20 | 750 | 270 | 730 | 650 |
| 21 | 760 | 270 | 720 | 650 |
| 22 | 770 | 270 | 710 | 650 |
| 23 | 790 | 270 | 690 | 650 |
| 24 | 800 | 280 | 680 | 640 |
| 25 | 820 | 280 | 660 | 640 |
| 26 | 830 | 290 | 650 | 630 |
| 27 | 850 | 300 | 630 | 620 |
| 28 | 860 | 310 | 620 | 610 |
| 29 | 880 | 320 | 600 | 600 |
| 30 | 890 | 340 | 590 | 580 |
| 31 | 900 | 350 | 580 | 570 |
| 32 | 910 | 370 | 570 | 550 |
| 33 | 920 | 380 | 560 | 540 |
| 34 | 920 | 400 | 560 | 520 |
| 35 | 930 | 410 | 550 | 510 |
| 36 | 930 | 430 | 550 | 490 |
| 37 | 930 | 440 | 550 | 480 |
| 38 | 930 | 450 | 550 | 470 |

**Table A2**. Overview of the number of trials included per participant and condition as well as the mean and standard deviation (sd) for each condition (bottom row). The maximum number of trials included was 38 per condition.

| Participant | Control | Empathy | Multi-motive | Reciprocity |
| --- | --- | --- | --- | --- |
| 1 | 21 | 37 | 38 | 38 |
| 2 | 28 | 28 | 36 | 25 |
| 3 | 19 | 18 | 14 | 13 |
| 4 | 24 | 21 | 28 | 26 |
| 5 | 36 | 37 | 38 | 37 |
| 6 | 33 | 32 | 33 | 32 |
| 7 | 22 | 26 | 24 | 26 |
| 8 | 21 | 23 | 23 | 17 |
| 9 | 23 | 19 | 22 | 23 |
| 10 | 30 | 31 | 29 | 33 |
| 11 | 32 | 27 | 33 | 36 |
| 12 | 16 | 20 | 19 | 16 |
| 13 | 19 | 20 | 20 | 19 |
| 14 | 38 | 38 | 38 | 38 |
| 15 | 36 | 38 | 38 | 18 |
| 16 | 33 | 32 | 29 | 34 |
| 17 | 28 | 27 | 36 | 37 |
| 18 | 14 | 24 | 26 | 23 |
| 19 | 32 | 30 | 27 | 29 |
| 20 | 21 | 24 | 20 | 21 |
| 21 | 19 | 18 | 20 | 18 |
| 22 | 31 | 33 | 31 | 31 |
| 23 | 15 | 34 | 31 | 32 |
| 24 | 36 | 35 | 31 | 29 |
| 25 | 19 | 34 | 27 | 30 |
| 26 | 20 | 34 | 27 | 28 |
| 27 | 25 | 32 | 31 | 21 |
| 28 | 6 | 36 | 38 | 31 |
| 29 | 33 | 34 | 37 | 33 |
| 30 | 37 | 34 | 38 | 36 |
| 31 | 33 | 36 | 37 | 30 |
| 32 | 23 | 31 | 30 | 30 |
| 33 | 26 | 23 | 22 | 26 |
| mean (sd) | 25.7 (7.9) | 29.3 (6.4) | 27.8 (7.0) | 29.4 (6.9) |

**Table A3**. Fixed effects results of the linear mixed model regression with the frequency of prosocial choices as dependent variable, induction rating, motive condition (empathy, reciprocity, multi-motive), and their interaction as fixed effects and participant as random intercept. Effects are reported with the empathy condition as reference level.

| Fixed effects | Beta | SE | χ^2^ | P |
| --- | --- | --- | --- | --- |
| induction rating | -0.046 | 0.118 | 6.380 | 0.012 |
| condition  multi-motive | -0.505 | 0.110 | 2.259 | .323 |
| reciprocity | -0.209 | 0.110 |  |  |
| induction rating*condition  multi-motive | 0.262 | 0.158 | 3.612 | 0.164 |
| reciprocity | 0.153 | 0.138 |  |  |

**Table A4**. Fixed effects results of the linear mixed model regression with the frequency of prosocial choices as dependent variable, induction rating, single motive condition (empathy, reciprocity), and their interaction as fixed effects and participant as random intercept. Effects are reported with the empathy condition as reference level.

| Fixed effects | Beta | SE | χ^2^ | P |
| --- | --- | --- | --- | --- |
| induction rating | 0.017 | 0.023 | 4.772 | 0.029 |
| condition  reciprocity | -0.048 | 0.042 | 0.156 | 0.693 |
| induction rating*condition  reciprocity | 0.035 | 0.025 | 2.055 | 0.151 |

**Table A5**. Fixed effects results of the linear mixed model regression with the frequency of prosocial choices as dependent variable, trait measure type (empathy, reciprocity), trait measure value (empathy trait value, reciprocity trait values), motive condition (empathy, reciprocity, multi-motive), and their interactions as fixed effects and participant as random intercept. Betas are reported with trait empathy and empathy condition as reference levels.

| Fixed effects | Beta | SE | χ^2^ | P |
| --- | --- | --- | --- | --- |
| trait_measure_value | 0.068 | 0.173 | 0.398 | 0.528 |
| trait_measure_type | 0.035 | 0.094 | 0.000 | 1.000 |
| condition  multi-motive | 0.022 | 0.110 | 12.381 | 0.002** |
| reciprocity | -0.226 | 0.110 |  |  |
| trait_measure_value*trait_measure_type  trait reciprocity | -.0148 | 0.120 | 0.000 | 0.986 |
| trait_measure_value*condition  multi-motive | 0.112 | 0.112 | 3.596 | 0.166 |
| reciprocity | -0.114 | 0.112 |  |  |
| trait_measure_type*condition  multi-motive | 0.000 | 0.156 | 0.000 | 1.000 |
| reciprocity | 0.000 | 0.156 |  |  |
| trait_measure_value*trait_measure_type*condition  trait reciprocity, multi-motive | 0.077 | 0.158 | 6.084 | 0.047* |
| trait reciprocity, reciprocity | 0.370 | 0.158 |  |  |

**Table A6**. Results of the pairwise Kolmogorov-Smirnov tests for the frequency of prosocial choices. Distance values D are given above the diagonal and p-values P below the diagonal.

|  | Baseline | Empathy | Reciprocity | Multi-motive |
| --- | --- | --- | --- | --- |
| Baseline | - | D = 0.21 | D = 0.18 | D = 0.24 |
| Empathy | P = 0.45 | - | D = 0.15 | D = 0.12 |
| Reciprocity | P = 0.65 | P = 0.84 | - | D = 0.12 |
| Multi-motive | P = 0.29 | P = 0.97 | P = 0.97 | - |

**Table A7**. Fixed effects results of the logistic mixed model regression with participants’s responses (prosocial vs. egoistic) as dependent variable, other possible gain, condition (baseline, empathy, reciprocity, multi-motive), and their interaction as fixed effects and participant as random intercept. Effects are reported with the baseline condition as reference level.

| Fixed effects | Beta | SE | χ^2^ | P |
| --- | --- | --- | --- | --- |
| other possible gain | 1.031 | 0.076 | 668.644 | <.001 |
| condition  empathy | 0.695 | 0.116 | 56.992 | <.001 |
| reciprocity | 0.363 | 0.111 |  | . |
| multi-motive | 0.763 | 0.118 |  |  |
| other possible gain*condition  empathy | 0.081 | 0.111 | 0.864 | .835 |
| reciprocity | 0.076 | 0.108 |  |  |
| multi-motive | 0.088 | 0.113 |  |  |

**Table A8**. Fixed effects results of the linear mixed model regression with reaction time as dependent variable, condition (baseline, empathy, reciprocity, multi-motive) as fixed effect and participant as random intercept. Effects are reported with the baseline condition as reference level.

| Fixed effects | Beta | SE | χ^2^ | P |
| --- | --- | --- | --- | --- |
| condition  empathy | -0.178 | 0.035 | 27.893 | <.001 |
| multi-motive | -0.128 | 0.035 |  | . |
| reciprocity | -0.131 | 0.036 |  |  |

**Table A9**. Points of subjective equality (PSE) for the different conditions and regression results. Mean values as well as the infimum (PSE – inf) and the supremum (PSE – sup) of the respective PSE are reported. Additionally, fixed effects results of the linear mixed model regression with participants’ PSEs as dependent variable, condition as fixed effect and participant as random intercept are reported with the baseline condition as reference level.

|  | PSE | PSE - inf | PSE-sup | Beta | SE | χ^2^ | P |
| --- | --- | --- | --- | --- | --- | --- | --- |
| condition |  |  |  |  |  | 2.888 | .409 |
| baseline | 107.17 | 81.70 | 127.27 |  |  |  |  |
| empathy | 41.81 | 6.18 | 69.97 | 0.324 | 0.239 |  |  |
| reciprocity | 76.56 | 47.81 | 105.89 | 0.322 | 0.239 |  |  |
| multi-motive | 32.29 | -8.19 | 59.49 | 0.347 | 0.239 |  |  |

**Table A10**. Fixed effects results of the linear mixed model regression with the frequency of prosocial choices as dependent variable, point equality, condition (baseline, empathy, reciprocity, multi-motive), and their interaction as fixed effects and participant as random intercept. Effects are reported with the baseline condition as reference level.

| Fixed effects | Beta | SE | χ^2^ | P |
| --- | --- | --- | --- | --- |
| point equality | 0.304 | 0.067 | 65.873 | <.001 |
| condition  empathy | 0.544 | 0.100 | 46.912 | <.001 |
| reciprocity | 0.279 | 0.097 |  |  |
| multi-motive | 0.601 | 0.101 |  |  |
| point equality*condition  empathy | -0.041 | 0.098 | 0.186 | 0.980 |
| Reciprocity | -0.014 | 0.096 |  |  |
| multi-motive | -0.025 | 0.099 |  |  |

**Table A11**. Overview of the models estimated and their DIC (deviance information criterion) values. The winning model is highlighted in bold font (the most complex model allowing all three parameters of interest to vary by condition). 1 = simple model, 2 = v varies by condition, 3 = z varies by condition, 4 = a varies by condition, 5 = v and z vary by condition, 6 = v and a vary by condition, 7 = z and a vary by condition, 8a = v, z, and a vary by condition, 8b = v, z, and a vary by condition and other gain is excluded as regressor.

| Model | Formula | DIC |
| --- | --- | --- |
| 1 | v ~ other gain | 11760.06 |
| 2 | v ~ other gain + condition | 11256.29 |
| 3 | v ~ other gain, z ~ condition | 11269.15 |
| 4 | v ~ other gain, a ~ condition | 11520.63 |
| 5 | v ~ other gain + condition, z ~ condition | 11174.30 |
| 6 | v ~ other gain + condition, a ~ condition | 11187.75 |
| 7 | v ~ other gain, z ~condition, a ~ condition | 11270.45 |
| 8a | v ~ other gain + condition, z ~ condition, a ~ condition | **11174.22** |
| 8b | v ~ condition, z ~ condition, a ~ condition | 12683.28 |

Table A12. Quantile comparison of the observed reaction time data with reaction time data simulated based on the drift diffusion model (500 simulations), as well as the standard deviation (std), standard error of means (SEM) and mean squared error (MSE) of the simulated data. The column “credible” indicates whether the data fall within the 95 % credible interval (if “True” the model is a 95% credible fit for the observed data). *ub* = upper boundary, *lb* = lower boundary.

| statistic | observed data | mean (simulated data) | Std | SEM | MSE | credible |
| --- | --- | --- | --- | --- | --- | --- |
| accuracy | 0.7497 | 0.7626 | 0.1897 | 0.0002 | 0.0362 | True |
| mean (ub) | 1.5968 | 1.6528 | 0.3819 | 0.0031 | 0.1490 | True |
| std (ub) | 0.6562 | 0.6446 | 0.3094 | 0.0001 | 0.0958 | True |
| 10q (ub) | 0.8721 | 1.0712 | 0.2571 | 0.0396 | 0.1057 | True |
| 30q (ub) | 1.1543 | 1.2856 | 0.2785 | 0.0172 | 0.0948 | True |
| 50q (ub) | 1.4700 | 1.4847 | 0.3182 | 0.0002 | 0.1015 | True |
| 70q (ub) | 1.8441 | 1.7637 | 0.4156 | 0.0065 | 0.1792 | True |
| 90q (ub) | 2.5369 | 2.3961 | 0.6991 | 0.0198 | 0.5086 | True |
| mean (lb) | -2.0119 | -2.1296 | 0.7201 | 0.0138 | 0.5323 | True |
| std (lb) | 0.6889 | 0.6756 | 0.4897 | 0.0002 | 0.2400 | True |
| 10q (lb) | 1.1550 | 1.4998 | 0.5995 | 0.1189 | 0.4783 | True |
| 30q (lb) | 1.5875 | 1.7348 | 0.6256 | 0.0217 | 0.4131 | True |
| 50q (lb) | 1.9645 | 1.9775 | 0.6929 | 0.0002 | 0.4803 | True |
| 70q (lb) | 2.3345 | 2.3089 | 0.8205 | 0.0007 | 0.6738 | True |
| 90q (lb) | 2.9565 | 2.8964 | 1.1242 | 0.0036 | 1.2674 | True |

**Table A13.** Relative differences between the baseline condition and the motive conditions for each of the three drift-diffusion modelling parameters of interest.

| Parameter | $\frac{\text{empathy-baseline}}{\text{baseline}}$ | $\frac{\text{reciprocity-baseline}}{\text{baseline}}$ | $\frac{\text{multi-motive-baseline}}{\text{baseline}}$ |
| --- | --- | --- | --- |
| *a* | - 1.64 % | -2.70 % | 3.33 % |
| *v* | -2.28 % | -0.835 % | -2.51 % |
| *Z* | 5.60 % | 1.65 % | 7.80 % |

**Table A14.** Neural results of the second-level regression between prosocial choice-related activity in the multi-motive condition > reciprocity condition and increase in prosocial choice preferences in the multi-motive condition relative to reciprocity (Δ*z*_multi-motive/reciprocity_) (P < .001 uncorrected, k > 10 voxels).

| Region | Hemisphere | x y z | Cluster size | *t*-value | P(FWE_cluster-corrected_) |
| --- | --- | --- | --- | --- | --- |
| Putamen | Right | 30 2 -2 | 143 | 5.49 | .001 |
|  | Left | -28 -9 1 | 62 | 5.36 | .003 |
| Middle cingulate gyrus | Right | 8 -24 31 | 29 | 5.73 | .442 |
| Posterior cingulate gyrus | Right | 8 -39 21 | 22 | 4.18 | .639 |
| Precentral gyrus | Right | 60 2 13 | 13 | 3.94 | .900 |
|  | Right | 43 -11 38 | 13 | 3.89 | .900 |
|  | Left | -58 -1 16 | 29 | 4.46 | .442 |
| Hippocampus | Right | 25 -9 -12 | 21 | 4.32 | .670 |
|  | Left | -30 -24 -15 | 12 | 4.43 | .922 |
| Insula | Left | -33 -11 18 | 21 | 4.39 | .670 |

**Table A15.** Results of the second-level one sample t-test of parametric modulation of neural activation by the partner’s gain during the prosocial decision process (p < .001 uncorr., k > 10 voxels)

| Region | Hemisphere | x y z | Cluster size | *t*-value | P(FWE_cluster-corrected_) |
| --- | --- | --- | --- | --- | --- |
| Insula | Right | 43 -6 18 | 108 | 6.52 | .009 |
|  | Left | -38 -916 | 517 | 6.23 | <.001 |
| Post central gyrus | Right | 60 -14 18 | 68 | 4.71 | .062 |
|  | Left | -40 -26 46 | 86 | 4.99 | .025 |
| Middle temporal gyrus | Right | 48 -56 18 | 75 | 4.82 | .044 |
| Pallidum | Right | 23 -9 -5 | 15 | 4.78 | .865 |
| Middle occipital gyrus | Right | 25 -89 13 | 85 | 4.67 | .027 |
|  | Left | -18 -94 13 | 22 | 4.16 | .676 |
| Fusiform gyrus | Right | 33 -74 -10 | 24 | 4.30 | .620 |
|  | Left | -30 -74 -12 | 16 | 3.88 | .841 |
| Middle cingulate gyrus | Left | -10 -24 48 | 11 | 4.02 | .945 |
| Superior temporal gyrus | Right | 58 -36 21 | 14 | 3.63 | .888 |
|  | Left | -50 -59 13 | 15 | 3.69 | .865 |
